# Root System Architecture and Environmental Flux Analysis in Mature Crops using 3D Root Mesocosms

**DOI:** 10.1101/2022.09.10.507424

**Authors:** Tyler G. Dowd, Mao Li, G. Cody Bagnall, Andrea Johnston, Christopher N. Topp

## Abstract

Current methods of root sampling typically only obtain small or incomplete sections of root systems and do not capture their true complexity. To facilitate the visualization and analysis of full-sized plant root systems in 3-dimensions, we developed customized mesocosm growth containers. While highly scalable, the design presented here uses an internal volume of 45 ft^3^ (1.27 m^3^), suitable for large crop and bioenergy grass root systems to grow largely unconstrained. Furthermore, they allow for the excavation and preservation of 3-dimensional RSA, and facilitate the collection of time-resolved subterranean environmental data. Sensor arrays monitoring matric potential, temperature and CO_2_ levels are buried in a grid formation at various depths to assess environmental fluxes at regular intervals. Methods of 3D data visualization of fluxes were developed to allow for comparison with root system architectural traits. Following harvest, the recovered root system can be digitally reconstructed in 3D through photogrammetry, which is an inexpensive method requiring only an appropriate studio space and a digital camera. We developed a pipeline to extract features from the 3D point clouds, or from derived skeletons that include point cloud voxel number as a proxy for biomass, total root system length, volume, depth, convex hull volume and solidity as a function of depth. Ground-truthing these features with biomass measurements from manually dissected root systems showed a high correlation. We evaluated switchgrass, maize, and sorghum root systems to highlight the capability for species wide comparisons. We focused on two switchgrass ecotypes, upland (VS16) and lowland (WBC3), in identical environments to demonstrate widely different root system architectures that may be indicative of core differences in their rhizoeconomic foraging strategies. Finally, we imposed a strong physiological water stress and manipulated the growth medium to demonstrate whole root system plasticity in response to environmental stimuli. Hence, these new “3D Root Mesocosms” and accompanying computational analysis provides a new paradigm for study of mature crop systems and the environmental fluxes that shape them.

## 1 Introduction

A plant’s root system is a complex set of organs that do more than simply anchor the plant to the ground and provide paths of uptake from the soil (Calvo et al., 2020; Novoplansky, 2019). Roots allow a plant to perceive its surroundings and adjust future growth accordingly, maximizing its chances of survival and reproduction (Bao et al., 2014; Dowd et al., 2019, 2020; Galvan-Ampudia and Testerink, 2011; Hématy et al., 2009; Knight, 1811; O’Brien et al., 2016). Thus, a plant’s Root System Architecture (RSA) is highly adaptable and is strongly affected by water and nutrient availability, competition with neighbors, rhizosphere interactions, and other aspects of the local growth environment (Gruber et al., 2013; Malamy, 2005; Morris et al., 2017; Rogers and Benfey, 2015; Yu et al., 2014). While it is widely accepted that understanding root form and function is one of the most critical aspects of plant biology, very little is known about below ground traits such as RSA compared to the wealth of information on above ground plant structures. As subterranean tissues with complex architectures that branch exponentially over time, they are very difficult to completely characterize, especially deep underground. Many methods exist to study root system architecture in various growth environments and growth stages (Atkinson et al., 2019; Dowd et al., 2021). However, all methods have significant tradeoffs, leading to the well-known gap between the information-dense data sets captured from plants grown in controlled environments, and the more realistic, but information-sparse nature of measurements collected from plants in the open field (Poorter et al., 2016; Topp et al., 2016).

Here we report the adaptation of traditional “root mesocosms” as a bridging system to facilitate the growth, excavation, and preservation of 3-dimensional (3D) RSA, while providing the unconstrained growth available in the field (Dowd et al., 2021; Odum, 1984). We incorporated sensor arrays to measure biologically relevant gradients and dynamics of environmental factors: matric (water) potential, temperature, and sub-soil CO_2_ content at various depths in the soil profile. We modeled the 3D environmental data to facilitate the comparison of the environmental conditions over time with the RSA, which in the future could be used for *post hoc* predictions of root activity and plasticity. Using photogrammetry (aka Structure from Motion, SfM), we generated highly detailed 3D reconstructions of the root systems and developed a pipeline for analysis across the soil profile. Accuracy of the 3D models was verified using manual ground truthing in 3D space. Clear differences among grass species RSA and in the effects of ecotypes and environments on RSA were measured as a demonstration of the flexibility and power of the approach.

## 2 Materials & Methods

### 2.1 Mesocosm construction and preparation

The mesocosm is composed of several subsystems: 1) The external frame, 2) the internal frame, and 3) the sensors. Additional equipment is needed to digitize, visualize and analyze the RSA information.

The external frame has a base constructed using pressure treated 10.2 × 10.2 cm dimensional lumber (i.e. 4×4s). The unit has a foot print of 135.3 cm x 109.9 cm. Four of the 4×4 pieces measuring 135.3 cm long are laid out parallel to each other, with one on each outside edge and two in the middle with a spacing of 10.8 cm. This configuration allows a standard pallet jack or forklift to pick up the unit. Two 4×4s measuring 109.9 cm are attached on top of the existing 4×4s using galvanized 1.9 cm bolts at each end, running perpendicular to create the rectangular base. Five 5.1 × 15.2 cm pressure treated yellow pine dimensional lumber boards (i.e. 2×6s) cut to a length of 109.9 cm were then laid out parallel to the top 4×4 boards, and attached to the first four 4×4 boards using 7.6 cm construction screws, thus creating a base for the mesocosm unit (Fig 1A). Four 4×4s that were 182.9 cm in length were inserted with a vertical orientation at the inside of the four-perimeter base frame. These vertically oriented 4×4s were attached using two 1.9 cm galvanized bolts. A drain box was constructed using 1.9 cm thick plywood that was constructed with an interior dimension of 91.5 cm x 91.5 cm (Fig 1B). The drain box was centered on the external frame base between the four vertical 4×4s. This box was lined with a polyvinyl pond liner and fitted with a 1.9 cm diameter drain pipe which stuck out the front of the mesocosm unit. The box was then filled with stones ranging from one to 7.6 cm diameter and an expanded metal top was placed on it.

**Figure 1:**
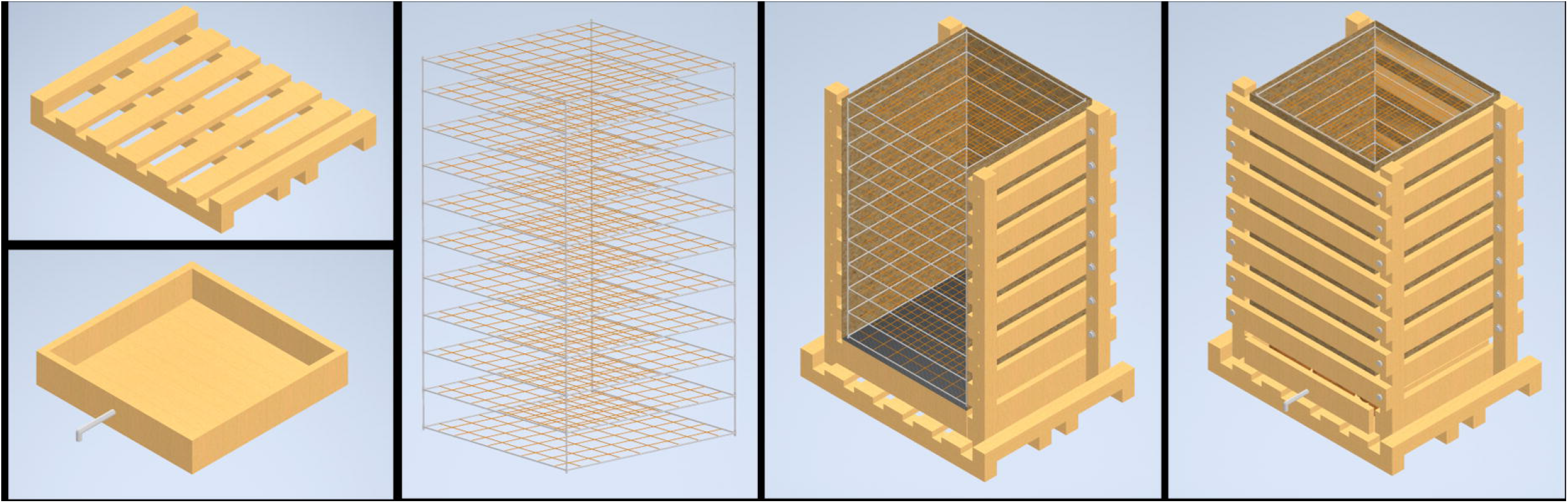
Major structural components of the mesocosm system. The mesocosm base is constructed from pressure treated lumber and designed for easy movement by a pallet jack (A). Directly above the base is a drainage box equipped with a drain spout to facilitate flow-through irrigation and allow sample collection (B). The internal component of the mesocosm is a scaffold system constructed of 0.5 inch PVC and fishing line (C). The exterior mesocosm walls are composed of lumber and held together with galvanized bolts (D). Between the internal frame and external lumber are thin boundaries of masonite that form smooth interior surfaces and a tarp to hold water in. The boards on the outside of the mesocosm can be attached and removed for easy access to the interior scaffold after roots have been grown.

The internal frame is used to support the roots, and maintain their spatial configuration when the roots are removed from the unit at the end of the experiment. The internal frame is constructed using a 1.3 cm nominal diameter polyvinyl chloride (PVC) pipe. The internal frame is a rectangular prism 152 cm tall consisting of 10 layers with each layer being 15.3 cm apart. Each layer of the frame is square in shape with a nominal length of 91.4 cm. Each side of the square has a 0.32 cm hole drilled at 10 cm increments. A polycarbonate line is strung across the frame connecting opposite holes, thus creating a 10 cm x 10 cm grid in the XY plane. When these squares are assembled together it creates a 10 cm x 10 cm x 15.3 cm grid in the XYZ planes (Fig 1C).

The drain pipe was used to designate the front of the unit. Eight 2×6 dimensional boards were used to connect the front vertical 4×4s on both the left and the right side. These 2×6s were attached on the inside face of the 4×4’s using 17.6 cm long construction screws, leaving 7.6 cm gaps between boards. The front and back of the unit had seven 2×6s connecting the left side to the right side. These boards were connected to the outside face of the 4×4 using galvanized 1.9 cm nuts and bolts, which allowed the boards to be taken on and off as needed (Fig 1D).

Once the external frame is constructed, four sheets of 0.3 cm thick particle board are cut to 121.9 cm long by 91.5 cm wide. These boards are placed with the long side in the vertical orientation, and on the inside of the external frame. A 16 mil (0.4 mm) thick polyethylene tarp is folded using an origami technique to create a rectangular prism shape that matches the external frame. The internal frame was then placed inside the tarp, and the front of the mesocosm unit was closed up (Fig 1E).fig

### 2.2 Environmental monitoring

#### 2.2.1 Matric potential, temperature and CO2 sensors

A variety of sensor arrays have been tested and deployed in the mesocosms system. Three metrics that have successfully been modeled to capture their dynamics in 3D space are the matric potential and temperature of the growth media as well as sub-soil CO_2_ levels. Temperature and matric potential are both measured via TEROS21 (Meter Group Inc., Pullman, WA, USA) sensors connected to Em50 data loggers (Meter Group Inc., Pullman, WA, USA) while CO_2_ measurements were taken using a Picarro G2201-i Isotopic Analyzer (Picarro Group, Santa Clara, CA, USA). By arranging the sensors in an array of 14 sampling points throughout the growth volume data interpolations allow the 3D modeling of the dynamic fluxes in the root system’s local growth environment (Fig 2). Matric potential and temperature measurements were set to record hourly, continuously. The CO_2_ profile throughout the growth volume was assessed by sampling air from rubber tubes buried in an array. CO_2_ measurements for each location in a mesocosm were sampled for 10 minutes and the mean value of the recorded CO_2_ levels were taken once weekly. Automation of sampling was facilitated using 14 ports on the Picarro 16-Port Distribution Manifold, set to switch through sample ports connected to each tube in the array.

**Figure 2:**
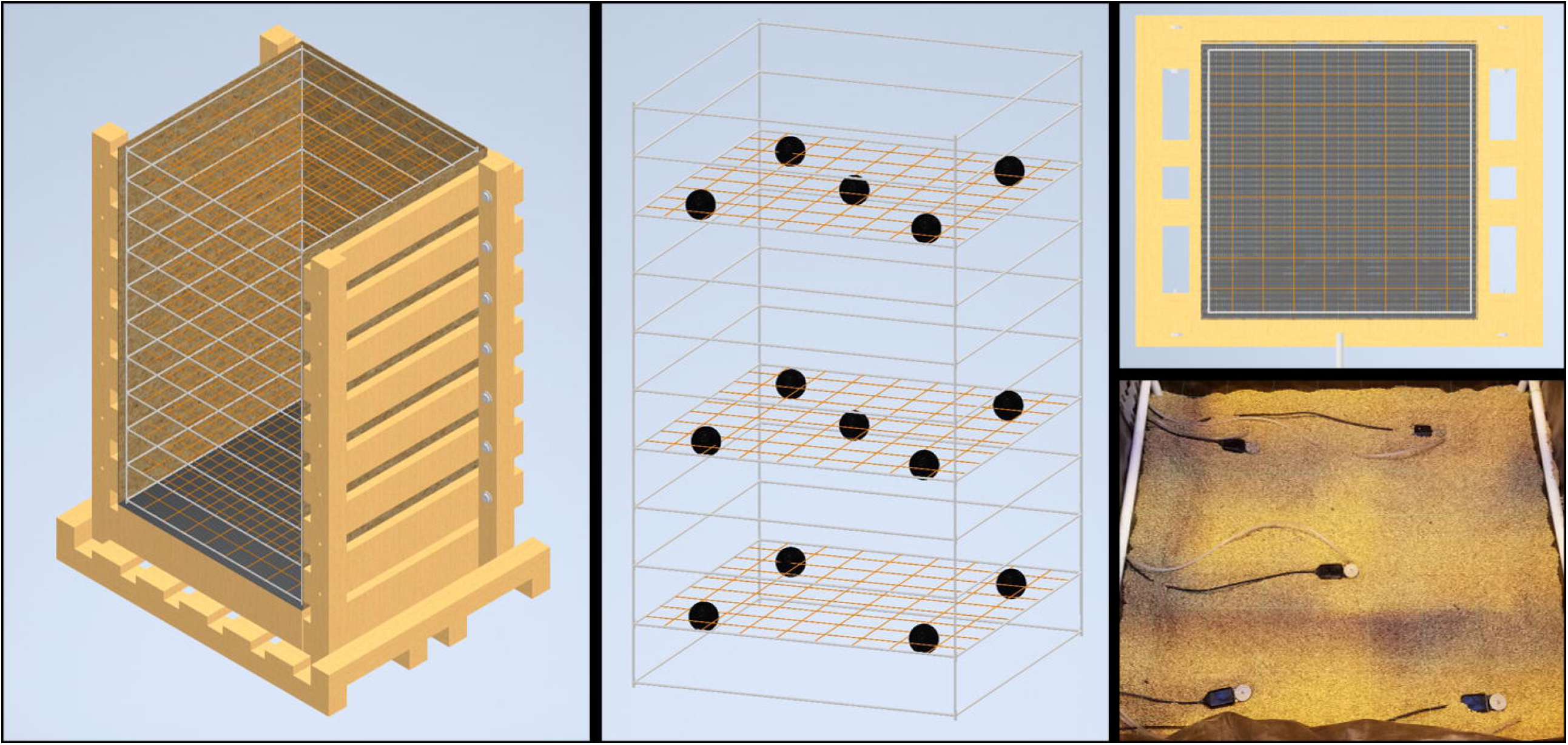
Interior sensor layout of the mesocosm. The interior PVC and fishing line scaffold create a coordinate system that can be used for sensor placement and data interpolation. Various environmental sensors (black spheres) were placed in a grid formation at 3 elevations throughout the mesocosm growth profile, 1.25 ft, 2.5 ft, and 4.25 ft deep. At the two upper elevations 5 sensors are laid out in a cross pattern while on the lowest level there are 4 with the center sensor absent. The lower right panel shows a photograph of both TEROS21 matric potential/ temperature sensors as well as air intake tubes for a Picarro gas analyzer.

#### 2.2.2 Data interpolation

We augment the 14 sensor data points (black points in Fig S1) to 35 data points by linearly calculating the data at the 21 additional locations on the boundary of the 3D mesocosm box (purple points in Fig S1). Note that we limit the maximum value of the water potential value to 0. These 21 boundary data points serve as boundary constraints for the 3D linear interpolation to the entire 3D mesocosm growth volume. This 3D linear interpolation is conducted by running the MATLAB function griddatan() which is a Delaunay triangulation based method.

### 2.3 Mesocosm Harvest

When the desired plant growth stage has been reached the mesocosms are prepared for harvest by shutting down all irrigation and removing all the associated components (Fig 3A). If the experimental design allows, it is beneficial to allow the mesocosms to dry for a few days before harvesting to ease growth media extraction. At this time all cables from sensors are disconnected from data loggers and the final data points are downloaded.

**Figure 3:**
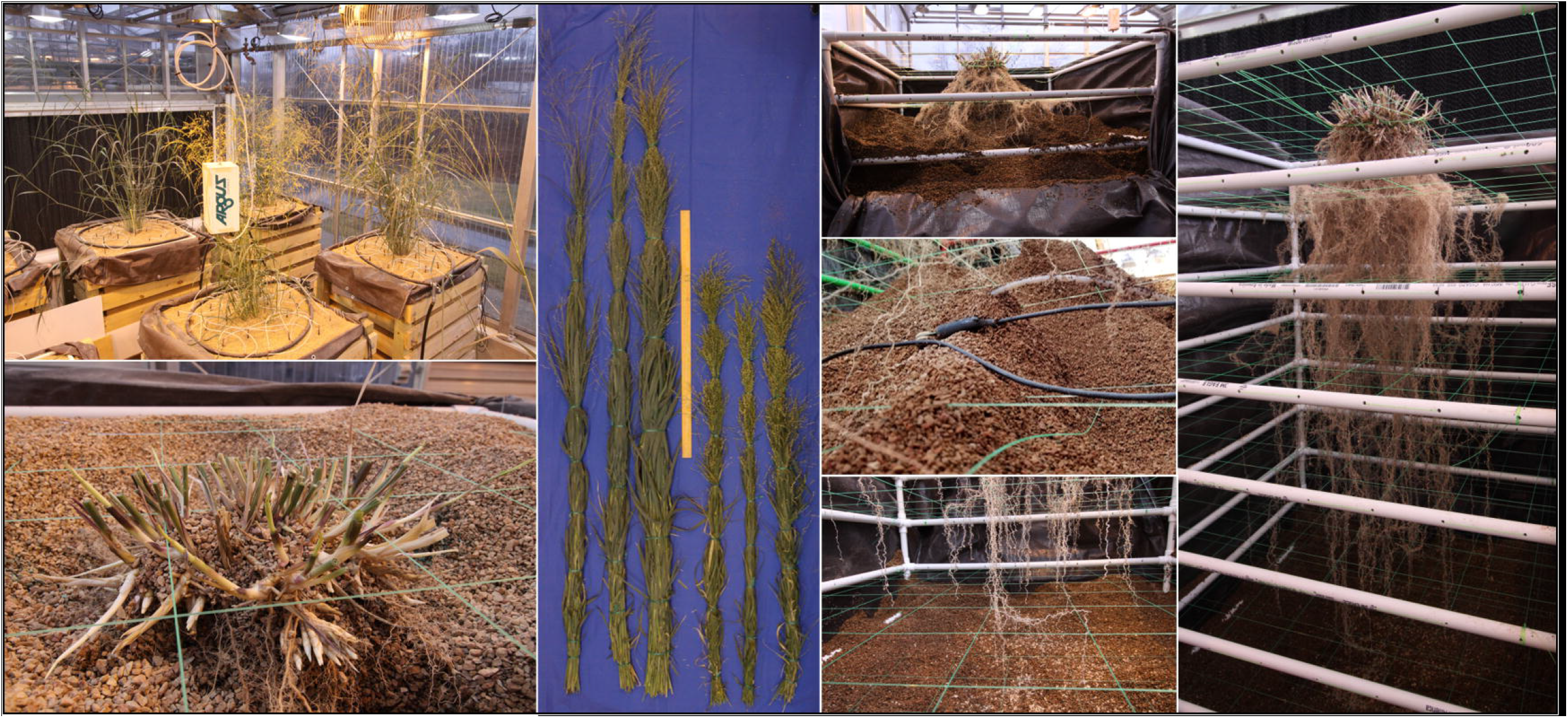
Mesocosm harvest method. Mesocosms supporting the growth of full size WBC3 and VS16 switchgrass genotypes (A). At harvest the shoots are cut a few centimeters above the soil profile (B). Shoots are bundled and dried for biomass measurements to accompany other shoot morphological traits collected during growth (C; Fig S2). Harvest begins by removing the uppermost exterior 2×6s, removing the masonite and pulling back the tarp to expose the top of the growth profile for excavation (D). Throughout sensors will be carefully extracted from the root system so as not to disturb the architecture (E). Harvest continues until all root tips are exposed from the growth media (F). Following complete excavation, the root system can be relocated in the PVC and fishing line scaffold and stored for future analysis (G).

At harvest, shoot tissues of the samples are harvested by cutting the plants near the surface of the growth media, above where the highest crown or adventurous root has emerged (Fig 3B). Shoot tissues are bundled together and are dried down to obtain biomass measurements (Fig 3C) to accompany any other shoot morphological data that was monitored during growth, such as plant height or tiller production (Fig S2).

In the absence of any shoot-born roots, it is still important to have a section of plant tissue above the growth media line to maintain proper orientation of the root system. Prior to the excavation of the root system the sample must be tied in place to maintain its position after the removal of the growth media. Additional fishing line, or other forms of support structures, can be used to tie the tissue emerging from the growth media surface (base of the shoot/ top of the root crown) to the top most section of the PVC frame. These supports go underneath the crown at the same height as the growth media and support the structure at the elevation it was at during growth. Tying the tissue off to all 4 sides will maintain the root crowns’ location in the X and Y orientations.

After securing the sample each of the eight 2×6 boards on the front and back of the mesocosm are loosened slightly to allow the removal of the particle board support on the front and back walls. Next the top pair of 2×6 boards are removed to expose the interior of the mesocosm and allow access to the uppermost layer of the growth media (Fig 3D). When excavating it is important to do so slowly as to not damage, or sever, unseen roots. During excavation, gentle vacuum suction is applied from the bottom of the exposed growth medium. This allows newly exposed roots to settle downward on the nearest segments of the interior scaffold to maintain root architecture. Caution must be taken to assure that the location of the vacuum tube is not in contact with any roots, direct suction can pull them from their location or cause them to snap. If rooting is too dense then manual hand clearing is necessary to excavate the section of the root system.

It is also important not to harvest too deeply in any given section as a shift in the growth media could lead to a landslide effect shearing roots in the process. This is more likely to happen if the growth media is wet and has high cohesion. Accordingly, the section of the growth media column that was exposed should be excavated completely before the next set of 2×6 boards are removed and the process repeats. If the mesocosm being harvested has sensors arrayed throughout the growth media, then each sensor is removed as they are excavated (Fig 3E). When the root tips of the deep axial roots are fully exposed, then less delicate methods of medium removal, such as handheld shovels, can be utilized to complete the excavation (Fig 3F).

After all of the growth media has been removed, the PVC frame can be slid out from the wooden exterior to provide 360 access to the exposed root system (Fig 3G). Depending on the growth medium used, an additional round of cleaning may be required to remove particles from dense areas of the root system. The now clean and free-standing root system can be stored for future analysis.

### 2.4 Photogrammetry

Utilizing 2D-photographs to develop a 3D point cloud through photogrammetry is a low-cost process requiring only a digital camera, an appropriate imaging studio, and photogrammetry software. Photogrammetry software identifies and utilizes a vast number of unique identification markers in each image to orient the photos in 3D space that share common markers. These can be natural/ architectural markers such as wood grain or lines between boards or bricks; or can be produced for the purpose of being a positional marker, such as painted shapes or specific computer-generated alignment patterns.

Our photogrammetry studio uses a combination of painted shapes (splatters and stencils), computer generated markers (code available on OpenCV: https://docs.opencv.org/4.x/d2/d1c/tutorial_multi_camera_main.html), and physical structures (AC unit, electrical control box, wire conduits, etc.) (Fig 4A). This studio has also been outfitted with many LED lights with very high color rendering indices and color temperatures of 5000K (daylight) to capture the most color accurate images possible. In the studio the sample is positioned in a central location between the lights to allow for full 360° movement around that sample and to minimize shadows (Fig 4B)

**Figure 4:**
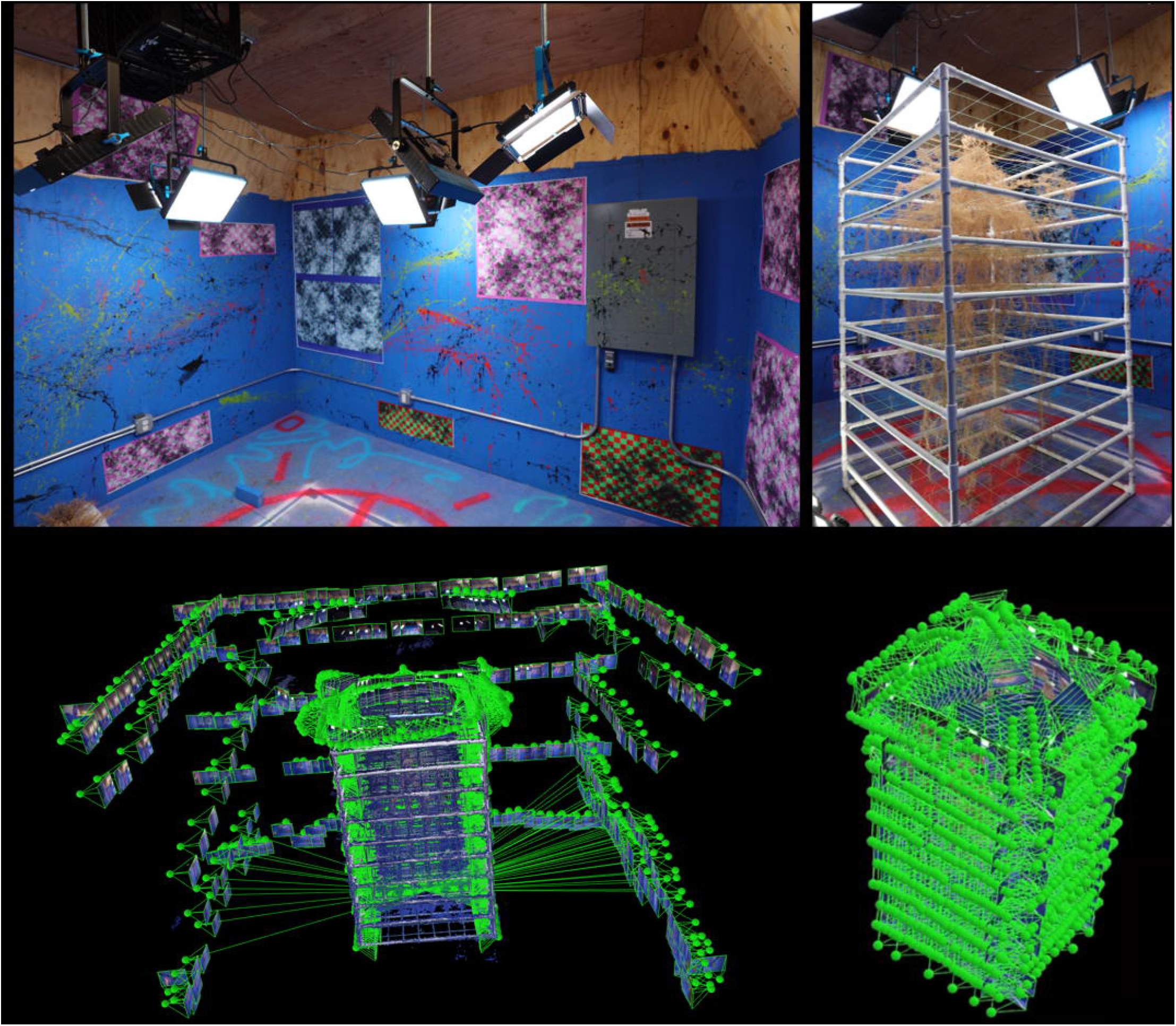
Photogrammetry studio and imaging process. Photogrammetry requires a dedicated space with many unique identification markers and strong uniform illumination (A). Plants samples are placed centrally in the studio to minimize shadows. Images are taken surrounding the subject (small rectangles are locations of individual camera locations) at several elevations to provide data to form the environment (C). Close up images are taken surrounding the root system with very high overlap to produce maximum system detail (D).

When imaging a sample by hand it is critical to use a sufficiently high shutter speed to ensure that the photographs of the sample and the environment remain in crisp focus. The images collected to produce the photogrammetry analyses detailed in this manuscript were taken at a shutter speed of 125 on a Canon EOS 50D (Ōta, Tokyo, Japan) set on the Tv (Time-value) priority setting. Ideally a fairly high aperture is also maintained to keep the entire root sample and identification markers within the depth of focus range. We found an aperture of 11-14 was ideal for the photogrammetry studio used in this study.

The zoom on the camera must not vary between images, as this will lead to artifacts in the resultant point cloud or failure of the photogrammetry software. Images were captured using a 10-18 mm wide angle lens with the zoom kept at 18 mm. It is important to keep the camera level during imaging which is monitored by an attached bubble level on the top of the camera.

It is critical that nothing is moved while the imaging is taking place. If an object in the environment (light plug cable, ground lights) or the sample itself is moved it will cause artifacts in the photogrammetry software. A small disturbance to the root scaffold will cause the very delicate roots to swing back and forth and it is likely that noise will be introduced into the point cloud. This could lead to a minor artifact, or possibly an entire doubling of the root system where two separate point clouds of the sample are produced with a slight offset.

The first images are taken along the perimeter of the photogrammetry room at a minimum of 4 elevations (eye level, chest, waist, and knees). This is to obtain a good baseline of the room and the ID markers (Fig 4C). This step will increase the match points of the up-close sample images and assist in camera alignment. Following this, images will be moved forward to be much closer to the plant sample. When imaging the sample up close photographs need to be captured on all sides as well as top-down images that angle smoothly from a dome shape to the flat walls of the root system scaffold. Images should have at least an 80% overlap (more is better) and should create a “dome” of coverage surrounding the sample (Fig 4D). A full-size crop plant root system is typically 4000+ images (including the images of the room perimeter).

Following image collection, the 2D-photographs are imported to a photogrammetry software to generate a 3D point cloud. The photogrammetry software found to perform the best with thin root structures is Pix4Dmapper (Pix4D S.A. Prilly, Switzerland). During the photogrammetric process voxels are mapped onto a 3D space to generate a 3D point cloud model of the sample. Once the point cloud is produced, many existing algorithms and processes utilized for X-ray CT or MRI data can be modified to analyze the point cloud and extract root system architectural traits.

### 2.5 Semiautomated segmentation of the root system point cloud

Following photogrammetry, the point cloud of the studio and surrounding environment is segmented away from the portion of the point cloud representing the root system. The 3D point cloud of the root system with the scaffold (white PVC pipes and green fishing lines) is loaded into Matlab (R2017a). Each point has its (x,y,z) coordinates and (R,G,B) color information. We segment out the root system from the point cloud by the following four steps.

#### 2.5.1 Linear transformation by aligning the scaffold point cloud to a predefined reference model

The first step standardizes the scaffold scale and position which is useful to remove the scaffold and extract features, especially the vertical distribution. We selected eight points from the 3D point cloud plotted in Matlab as target points. Four of these eight points are picked from the crossings of the fishing line grid on the top layer. The other four are chosen from the bottom layer (red points in Fig 5A). To be able to visualize and select the points more easily, we only work on the local layer containing the target points (middle panel in Fig 5A). The reference model is defined based on the scaffold design. Then the control points on the reference model are set (right panel in Fig 5A). Note these eight target points can be arbitrarily selected as long as they are not on the same plane and the control points correspond correctly. A Procrustes alignment is performed to determine a linear transformation (translation, rotation and reflection, scale) based on the target points and control points. We then apply these components to transform the entire 3D point cloud.

**Figure 5:**
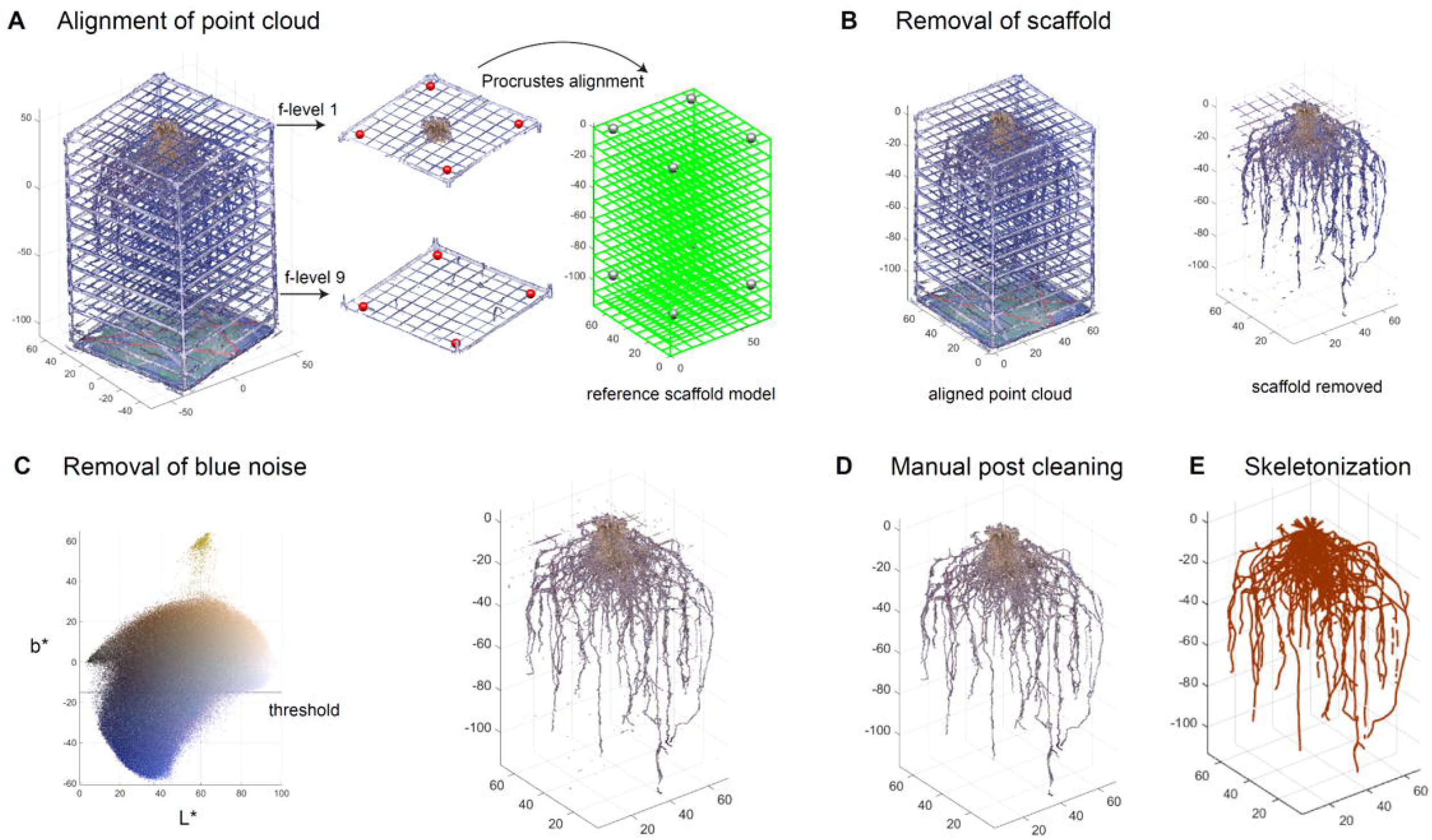
Semi-automated segmentation of the root system point cloud. Point cloud of the root system with the scaffold is aligned to a predefined reference model by performing linear transform on manually picked control points to the target points (A). Point cloud of the scaffold is removed based on the position and color information (B). Blue noise on the root is then removed using a threshold method (C). A post manual processing is conducted to further clean the root system point cloud (D). Point cloud is skeletonized into a network system using an algorithm based on a Laplacian contraction method (E).

#### 2.5.2 Removal of the scaffold

Although the scaffold now is aligned with the reference model (left panel in Fig 5B), we cannot simply delete the points along the reference as roots could be in contact with or be growing along the scaffold. Therefore, we determine the scaffold points not only based on the position, but also on the color. We set a small neighbor region near the reference model in case the candidate scaffold is slightly misaligned with the reference. We then define color thresholds to remove non-root points such as white, gray, and green points (right panel in Fig 5B).

#### 2.5.3 Removal of background noise

Additionally, it is likely that some blue color from the photo studio background could be merged into the point cloud during the 3D reconstruction. We would like to remove the blue noise. We convert the RGB color into CIELAB (L*a*b*) color in which L* represents lightness, a* represents green to magenta, b* represents blue to yellow. We used a practical threshold (b* =15) which separates the blue noise with the root (left panel in Fig 5C). The output of this process is a point cloud devoid of artifactual color noise and natural in appearance (right panel in Fig 5C). However, it may still contain some noise for various reasons, such as light refraction through the translucent fishing line giving it a color similar to the surrounding roots. At this point the root system point cloud is saved as a .ply for the manual post-cleaning process.

#### 2.5.4 Manual post-process cleaning

Once the point cloud has gone through segmentation in MATLAB, the data is further cleaned to remove unwanted artifacts, such as residual fishing line and noise. We make these changes in CloudCompare (v2.11.1 (Anoia), 2022) where the image can be cleaned using precise segmentation. Post-process manual cleaning allows for better accuracy of the root structure and can drastically improve the clarity of the 3D root system model (Fig 5D).

Noise on the point cloud at this stage is common, such as artifactual points in a cloud system that are not in the proximity of other roots, or remaining color transferred from the studio environment that was not completely removed by the color thresholding. Furthermore, due to the structural methodology of the mesocosm, it is necessary to remove certain artifacts from the point cloud that remain after segmentation, such as remnants of the PVC and fishing line scaffold. The segmentation tool is used to remove the noise and remnants, leaving an isolated root system (Fig S3).

Additionally, some areas of the point cloud will need manual correction and shaping. This is utilized predominantly in locations where tape has been placed to keep the roots together if they have broken during harvest or storage. The taped area appears larger and a different color in the point cloud but can be shaved down using precise segmentation. Shaping and smoothing can also eliminate areas of noise or unwanted artifacts. Once all artifacts are removed, the point cloud can be used for trait extraction and skeletonization (Fig 5E).

### 2.6 Root trait extraction from point clouds

From the point cloud, we can directly measure some global traits such as total number of points, convex hull volume (the volume of the smallest convex set containing the point cloud), elongation (PCA on point cloud, taking the ratio between PC2 variance and PC1 variance), flatness (the ratio between PC3 variance and PC2 variance), and maximum depth (the depth of deepest root point). We also can measure the vertical distributions for biomass (Gaussian density estimator for point cloud), convex hull volume (Gaussian density estimator for point cloud extracted from convex hull area at each depth), and solidity (spline interpolation of solidity through every depth). These distributions are then discretized into 10 bins for downstream analysis.

However, point clouds are made of scattered points without connection information. Volume and length-related features cannot be directly measured. To be able to compute the volume dependent features, we compute alpha shapes with a set of radii to form a few bounding volumes that envelop the point cloud (Edelsbrunner et al., 1983). Intuitively, an alpha shape is formed by scooping out ice cream with a sphere spoon without bumping into chocolate pieces (the points) and then straightening the boundaries. The size of a spoon is a parameter denoted as alpha. We measure these alpha shape volumes with three different scales (alpha =0.5, 1, and 2) which indirectly describe the root volume (Fig 6). We calculate the solidity using the ratio between alpha shape volume at alpha =2 and convex hull volume. To be able to compute the length dependent features, point cloud is skeletonized into a network system using an algorithm based on a Laplacian contraction method (Cao et al., 2010), which was conducted in Matlab R2017a. Then we can calculate length dependent features such as the total root length.

**Figure 6:**
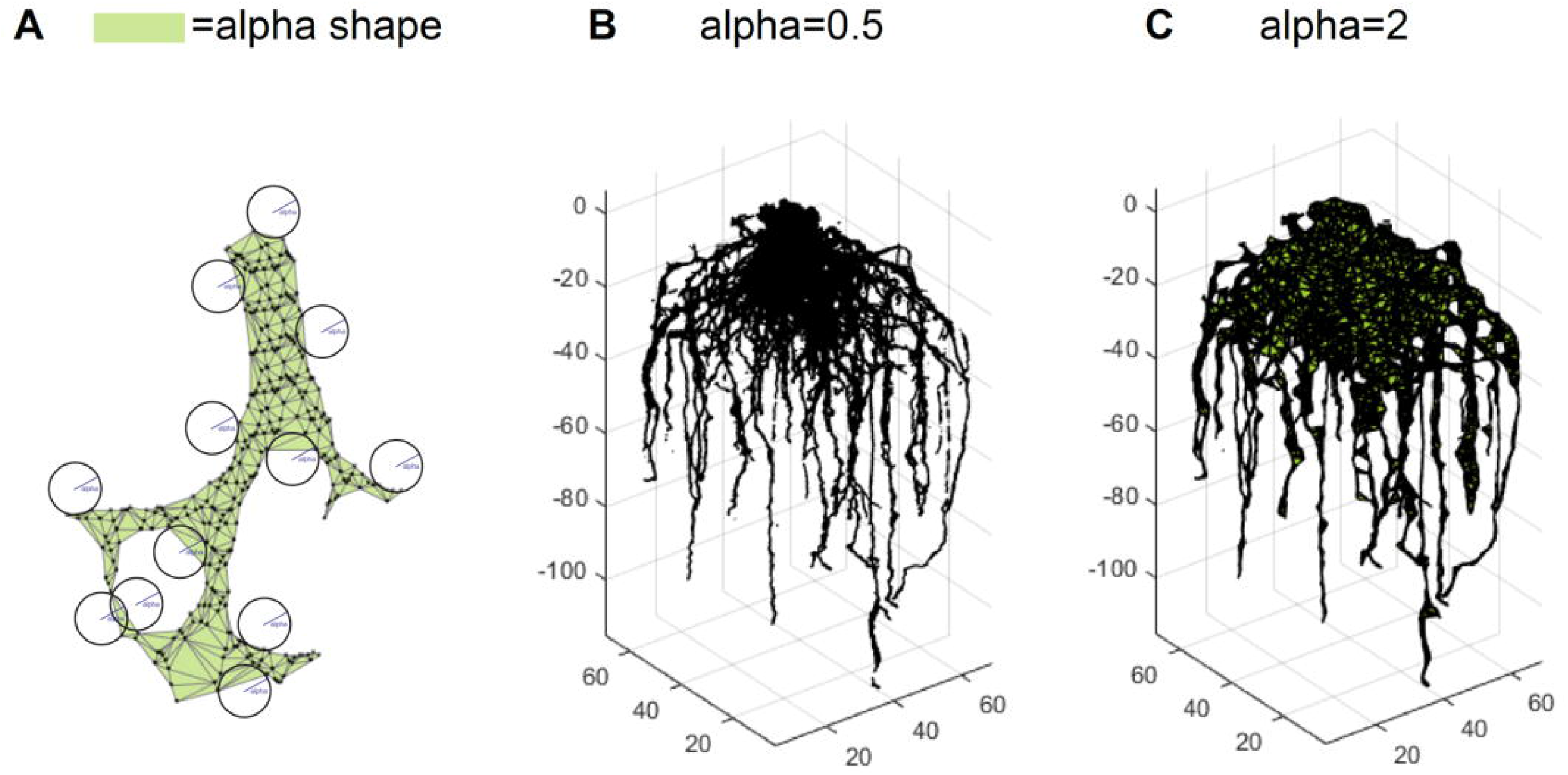
Trait extraction via alpha shape analysis. An alpha shape is a kind of shape that envelopes the point cloud (A). Intuitively, it is formed by scooping out ice cream with a sphere spoon without bumping into chocolate pieces (the points) and then straightening the boundaries. The volumes of alpha shape can be calculated for different parameters such as alpha=0.5 (B) and alpha=2 (C).

### 2.7 Root system 3D biomass

Biomass measurements are taken by utilizing a grid system. Each layer of the mesocosm, starting from the bottom, may contain biomass and is weighed. This process starts by identifying the location of the sample in the coordinates created by the fishing line structure. Mass is weighed by cutting the roots at each layer and recording the weight within each box. After completing each layer, the crown of the root is then removed, labeled, and stored for further analysis.

## 3 Results

### 3.1 Species and genotype modeling facilitated by the mesocosm systems

We successfully grew and modeled the entire root system architectures of mature (after flower formation) maize (PHZ51), sorghum (BTX623), and switchgrass (WBC3, VS16) (Fig 7; Supplemental Videos 1-4, Table 1) in Turface MVP (Profile Products LLC., Buffalo Grove, Ill) using our 3D Root Mesocosms. Variation in the root systems of these species is evident both by eye and through the analysis of the subsequent point clouds developed through photogrammetry. The bulk of our studies focused on two key switchgrass varieties that have adapted to different natural environments: upland (VS16) and lowland (WBC3) switchgrass (Milano et al., 2016). The distinct root system architectures of these genotypes are apparent (Fig 8A, D, G). While the upland VS16 genotype is smaller, it shows much less horizontal growth compared to the lowland WBC3, apparently prioritizing carbon allocation to deeper rooting under our experimental conditions. Furthermore, VS16 shows more vigorous lateral root growth relative to the total root system size and has a higher root to shoot ratio, responses believed to aid in capturing as much water as possible from the local environment. Conversely, WBC3 shows a much wider horizontal spread of water transporting axile roots coupled with less investment into water absorbing lateral roots, a pattern expected in plants adapted to environments with ample water availability (Weaver, 1926).

**Table 1:** Root-to-shoot ratios of various species and treatment combinations grown in 3D mesocosms Biomass data of dried shoot and roots weights, and the corresponding root-to-shoot ratios, for switchgrass (WBC3, VS16), sorghum, and maize grown in turface or mixed media under well-watered or water-stressed conditions. Data are means ± standard error, n= 3 for all treatments except Sorghum WS Turface (n=2) and Maize WW Turface (n=6).

|  | Shoot Weight (g) | Root Weight (g) | Root: Shoot Ratio |
| --- | --- | --- | --- |
| WBC3 WW Mixed Media | 265.3 ± 37.8 | 126.5 ± 26.9 | 0.48 |
| WBC3 WW Turfaced | 271.1 ± 79.5 | 144.9 ± 41.7 | 0.53 |
| WBC3 WS Turfaced | 44.5 ± 14.6 | 52.9 ± 14.3 | 1.19 |
| VS16 WW Mixed Media | 36.2 ± 16.4 | 22.0 ± 1.6 | 0.61 |
| VS16 WW Turfaced | 114.8 ± 23.3 | 98.6 ± 8.6 | 0.86 |
| Sorghum WW Turfaced | 257.6 ± 28.1 | 77.6 ± 2.0 | 0.30 |
| Sorghum WS Turfaced | 98.7 | 80.3 | 0.81 |
| Maize WW Turfaced | 75.9 ± 12.2 | 62.1 ± 8.4 | 0.87 |

**Figure 7:**
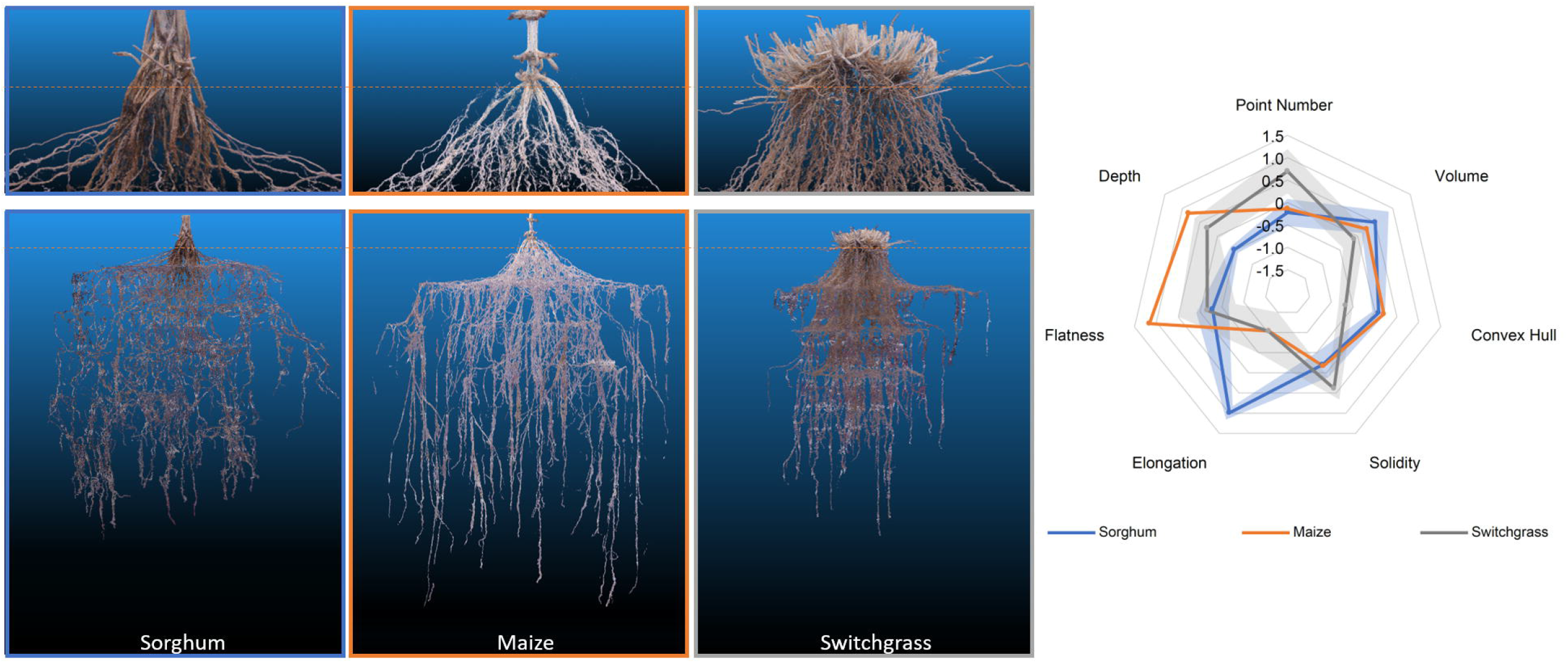
RSA of 3 different mesocosm grown species (left). Representative point clouds for sorghum, maize, and switchgrass species. Orange dotted line denotes the approximate growth media level during growth. A radar plot detailing the analysis of 7 different root shape traits from the point clouds. Data shown are mean ± standard error in shaded regions. Sorghum and switchgrass n=3, maize n=2.

**Figure 8:**
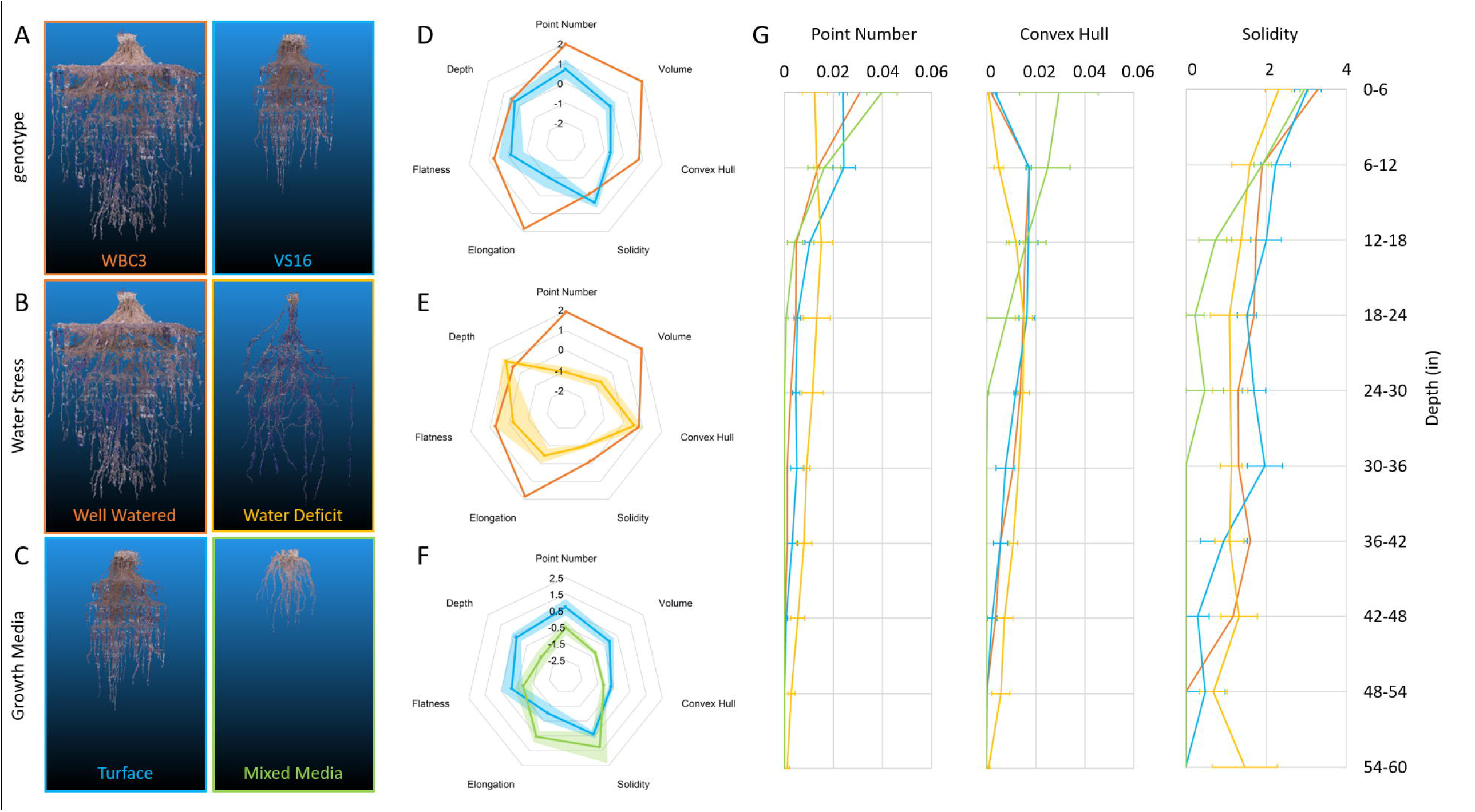
RSA traits of switchgrass affected by genetics and environmental conditions. Representative point clouds and extracted root traits of various G x E experimental conditions examinable via mesocosms. Genotypic comparison of WBC3 (orange) and VS16 (blue) when grown in well-watered turface (A, D). RSA response of WBC3 to well-watered (orange) and water stressed (yellow) turface conditions (B, E). RSA response of VS16 when grown in well-watered turface (blue) or a 3:1 potting mix to turface blend (green). Radar plots of data have all been standardized to allow comparison across treatments and traits, data shown are mean ± standard error. Point number (biomass proxy), convex hull, and solidity trait values (G) are presented for the entire depth of the growth media profile (WBC3 WW turface, orange; VS16 WW turface, blue; WBC3 WS turface, yellow; VS16 WW mixed media, green). Values for solidity were transformed by log(x*10000) for data visualization.

### 3.2 RSA model accuracy confirmed by 3D biomass ground truth

To ensure that the point clouds derived via photogrammetry are accurate to the actual RSA, a direct comparison to biomass in 3D space was necessary. Using the location of the internal mesocosm fishing line scaffold coordinates the root systems were dissected both physically and computationally (Fig 9, Fig S4). Using the 810 individual subunits formed by the scaffold the point cloud and biomass can be compared at a 10 cm x 10 cm x 15.25 cm resolution. Biomass ground truth measurements align well with *in silico* generated cubes of the point clouds that occupy the same space (Fig S5). Scaling the values of each coordinate section to the entirety of the growth space, the biomass and voxel amount, can be directly compared.

**Figure 9:**
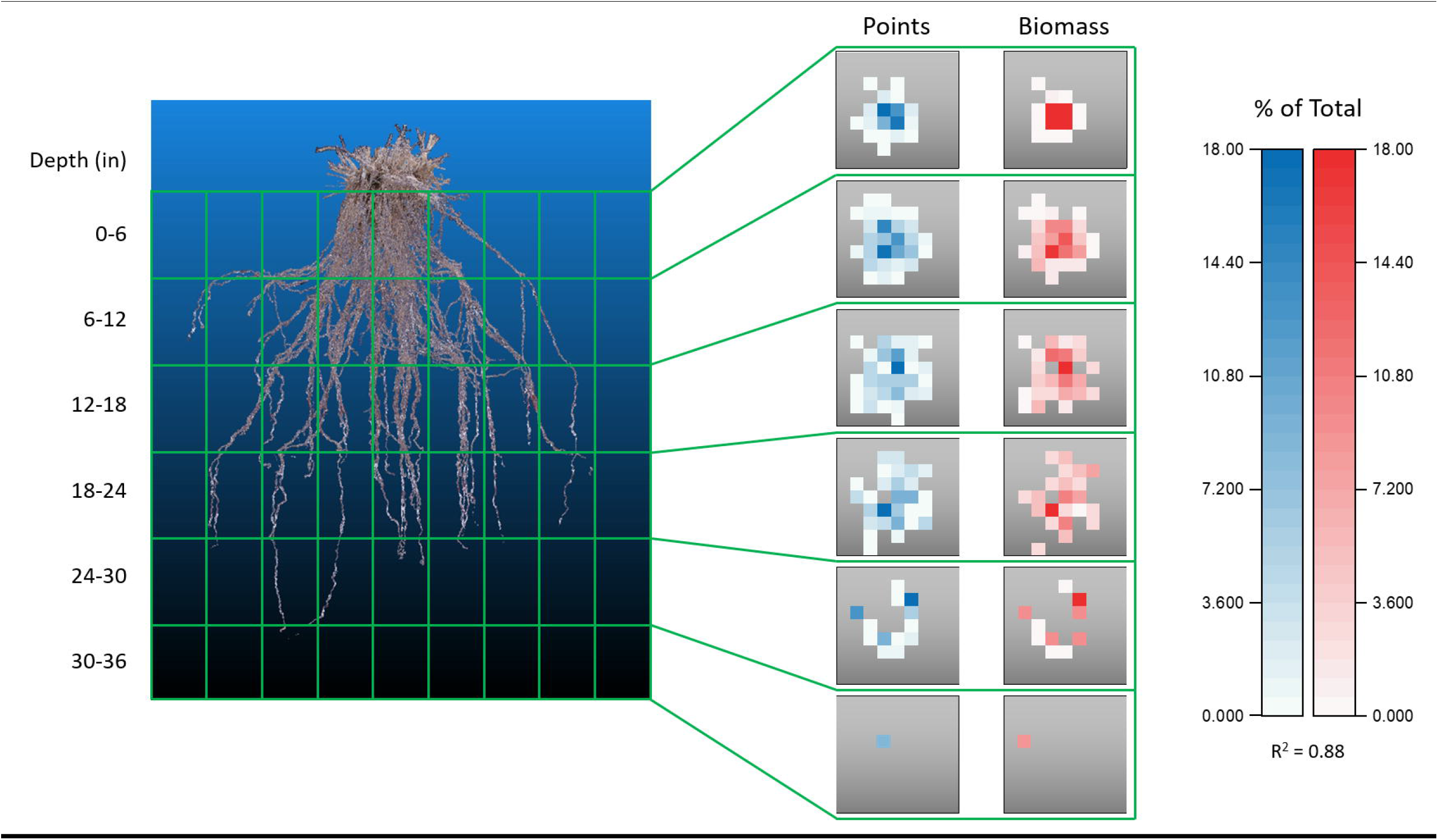
Biomass confirmation of point cloud accuracy. Comparison of data obtained from a switchgrass root system that had been physically and digitally dissected into the 180 independent sections outlined by the internal PVC and fishing line scaffold. Each gray square is a top-down view of a z-layer consisting of 9 × 9 cuboids. Data within each square represents the number of points, or the fraction of biomass, found in a cuboid as a percentage of the entire root system. Physical segmented biomass values correlate well with values of point number located in the same cuboid coordinate position when assessed on a relative scale, R^2^ = .088.

Beyond acting as a ground truth for point clouds, the biomass measurements obtained give an unprecedented sampling of entire root systems of full-grown crop plants largely preserved in their natural configurations. Out preliminary experiments show that differences can be observed between switchgrass genotypes, as well as in response to water stress (Fig 10). When grown under well-watered conditions both WBC3 and VS16 root systems displayed a similar profile of biomass allocation with depth, with the majority of biomass allocated in the upper profile and less allocated to each subsequent depth. In contrast, when WBC3 plants were grown under water stressed conditions the biomass allocation was modified and near-even amounts of root tissues were distributed at all depths down to 3 ft (91.44cm).

**Figure 10:**
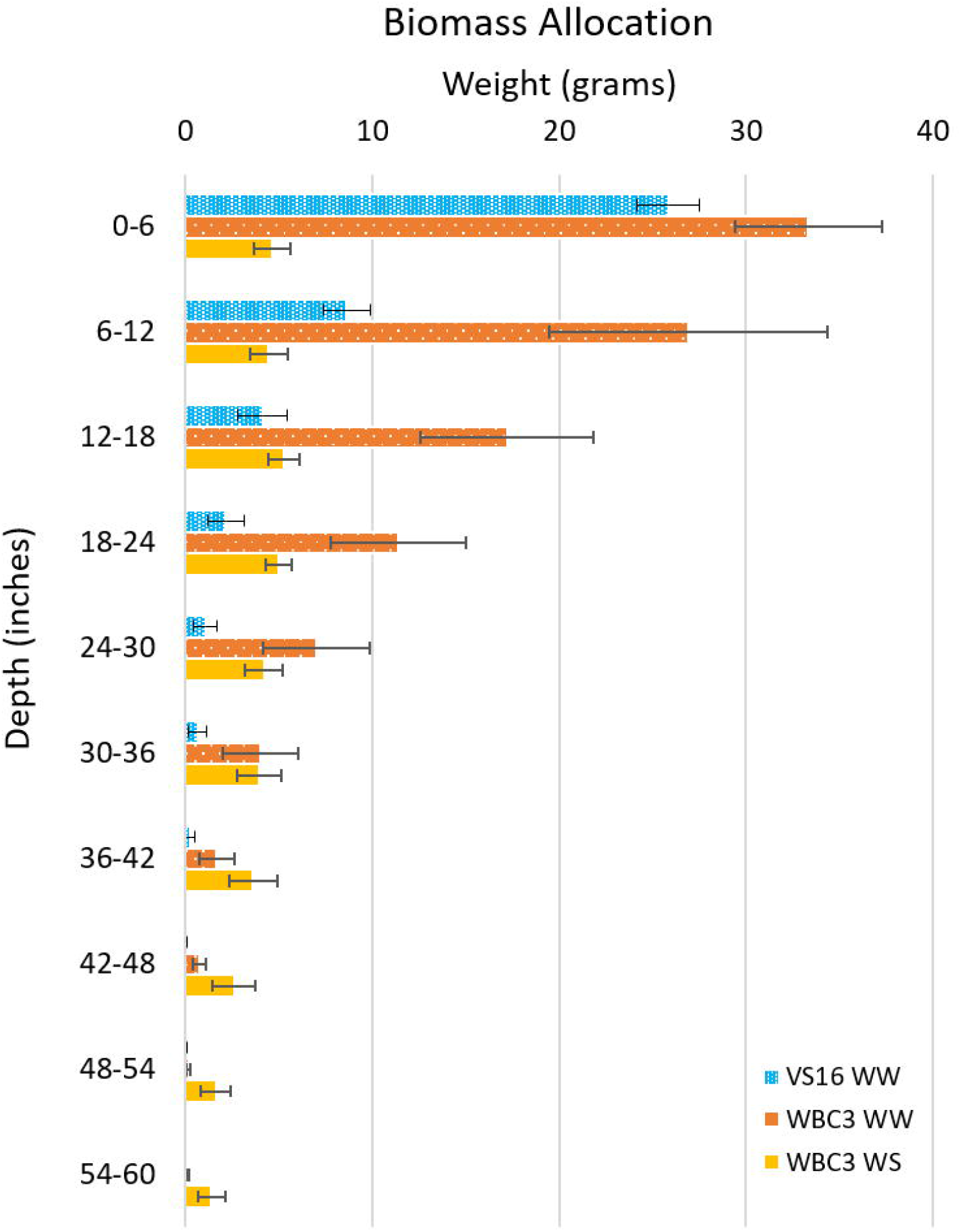
Biomass allocation by depth of switchgrass. Plot shows biomass measurements throughout the growth media profile for VS16 well-watered (blue), WBC3 well-watered (orange), and WBC3 water stressed (yellow). Data values are means ± standard error.

### 3.3 Mesocosms as a platform for water deficit experiments

The ready control and measurement of various environmental conditions in the 3D root mesocosms was demonstrated using a TEROS21 sensor array (Fig 2) to investigate the 3D root system phenotypic response of WBC-type switchgrass to physiologically-defined water stress. The high spatial and temporal resolution of our imputed sensor data (Fig S1) facilitated 4D monitoring of water fluxes from which we made delicate adjustments of irrigation to impose two levels of water availability: a well-watered treatment with a constant matric potential of -0.01 MPa and a water stress treatment with the average stress levels of approximately -2.5 MPa.

Continuously monitoring the matric potential revealed the real-time dynamics of water deficit throughout the duration of plant growth, including diurnal patterns of wetting and drying tied to daily transpiration (Fig S6). The TEROS21 system simultaneously collects temperature data which can be analyzed in conjunction at the same resolution (Fig S6). We note the temperature gradient in our system mimics field soils to an extent, insofar as temperature decreases with depth.

Several hallmarks of traditional responses to water deficit were seen in WBC3 when grown under the moderate-to-severe level of stress (-2.5 MPa), including a major reduction in root system volume and convex hull, but with maintenance of overall root system depth (Fig 8, B, E; Supplemental Video 5), leading to a significant shift of the root to shoot ratio (Table 1). The tradeoff to maintaining depth with a smaller root mass is a reduced global solidity, which quantifies the thoroughness of soil exploration in the rooting zone (defined by the convex hull volume). Analysis of root system traits across the depth profile revealed the biomass and convex hull area of water-stressed WBC3 was larger than well-watered below ∼12 inches (∼30cm), revealing allocation of more biomass (point number) to root proliferation at depth (Fig 8G). However, in the upper profile WBC3 displayed more biomass and a larger convex hull under well-watered conditions compared to water stressed, with ∼71% of the total root mass in the top ∼12 inches.

### 3.4 Assessing effects of growth media on RSA and the root zone environment

To study the effects of growth media on RSA and environmental parameters, we explored the incorporation of standard greenhouse potting mix (Berger BM7, Berger Saint-Modeste, QC) into the system under well-watered conditions (Fig 8C, F, G). When grown in a mix of 3:1 potting mix to turface WBC3 plants appeared to have longer and less branched lateral roots than when grown in pure turface (Fig S7). We suspect this change is a response to the particle size of the potting mix, which is much smaller than the average turface particle, and has a greater hydraulic conductivity. Roots growing though smaller potting mix particles require less lateral branching to access growth media bound water as there would be significantly more root-to-particle contact points along the root compared to growth in the turface. The VS16 plants developed very small root systems under the mixed media compared to the turface, as well as reduced root to shoot ratios, perhaps reflecting that they are not adapted to grow in an extremely wet environment (Fig 8 C, F, G; Table 1).

We used a different facet of our 3D sensor array to monitor dynamic CO2 respiration from soils (Fig 11). Both root and microbial respiration are major drivers of subsoil CO_2_ production, and rhizosphere processes such as microbial consumption of root exudates and soil organic matter link these pools. Turface is a calcined clay product and contains little or no organic matter (OM) (Beddes et al., 2013; Beddes and Kratsch, 2009; Calonje et al., 2010), whereas greenhouse potting mix typically has a very high OM content (in our case Berger BM7 is ∼79%). In turface at early time points, VS16 and WBC3 switchgrass CO_2_ profiles are very similar, although VS16 is set higher (Fig 11C, D). Over time (around week 8) CO2 levels at all three measured depths begin to rise, presumably as a result of rapid root proliferation. However, WBC quickly rises several-fold at the lowest depth (4.5 feet), consistent with differences in its eventual root system size (Fig 8A).

**Figure 11:**
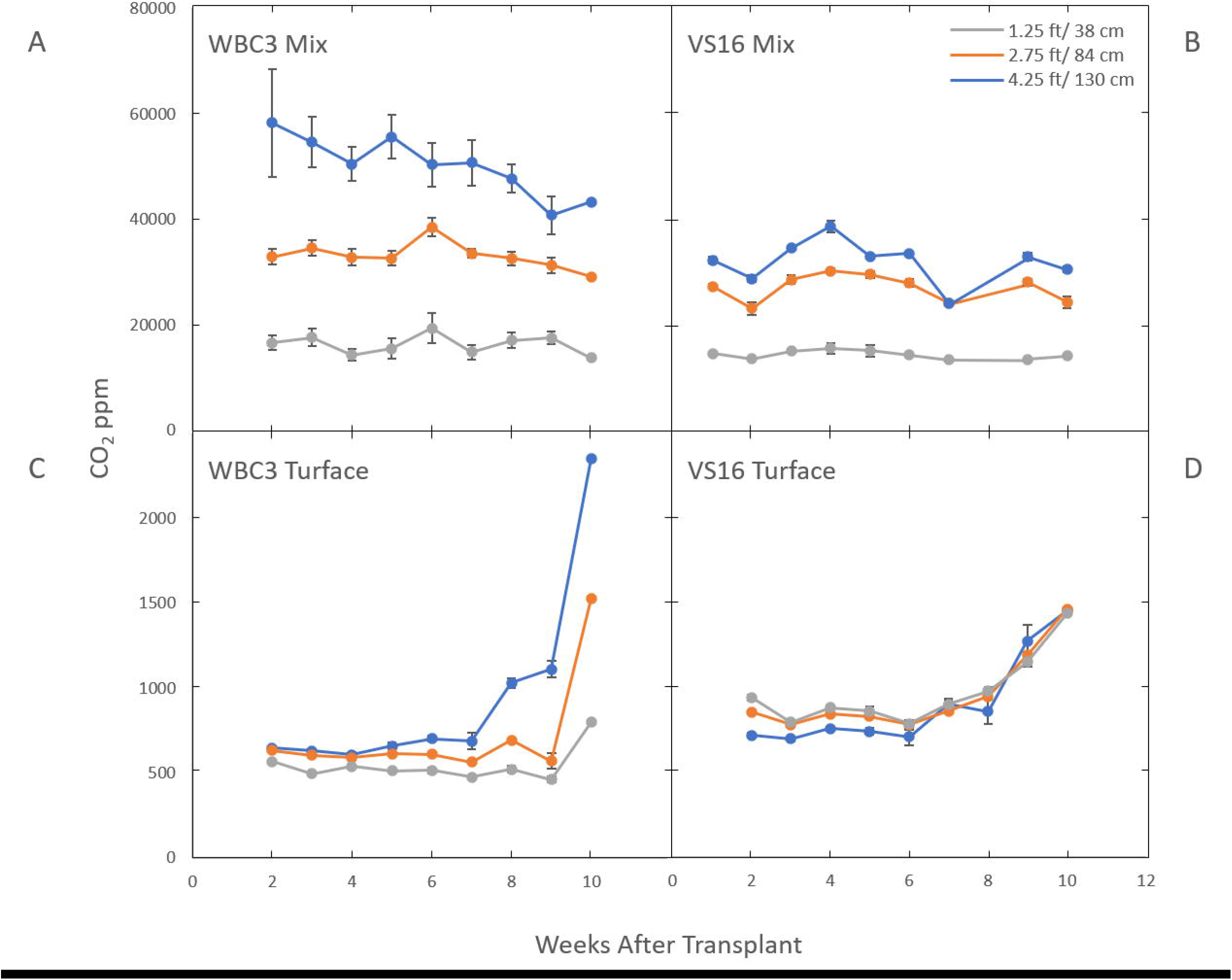
Subterranean CO_2_ levels in the mesocosms are affected by which switchgrass genotype is growing in what growth media. Graphs show CO_2_ levels in mesocosms growing WBC3 (A, C) and VS16 (B, D) grown in pure turface (C, D) and a 3:1 potting mix to turface blend (A, B). Measurements were taken at three depths, 1.25 ft (gray), 2.75 ft (orange) and 4.25 ft (blue) below the soil profile. Data values are means ± standard error.

Interestingly this same relationship is not seen in mesocosms filled with a 3:1 potting media: turface mix. Under these conditions the CO_2_ levels were several orders of magnitude higher compared to turface filled mesocosms and the levels remained constant or showed a slight decline throughout the growth term (Fig 11A, B; Fig S8). The significantly higher CO_2_ levels and their stability at the sample locations at the middle and higher elevations suggest that a combination of the organic components and microbial population of the potting mix play a more significant role than direct root respiration. Yet, WBC3 mesocosms showed elevated CO_2_ levels in the lowest growth media profile compared to VS16 mesocosms, an area where VS16 root systems did not occupy (Fig 8A). This result suggests that local root activity at depth in WBC samples may be driving increased microbial activity via rhizosphere priming (Kuzyakov, 2002) (Fig 11A, B).

### 3.5 Combined analysis of RSA models and 3D environmental data

Aligning the photogrammetry point clouds with the time course 3D environmental data fluxes provides the opportunity to make post hoc hypotheses on how the environment shaped the mature RSA. Alterations in matric potential and temperature in the growth media along the path of root development can give insight into the conditions that resulted in the RSA, and changes in sub soil CO_2_ are correlated with the presence of root respiration (Fig 12; Supplemental Videos 6, 7). This type of analysis can be used to make observations to provide training data to a model in an effort to estimate root location and activity based on localized environmental fluxes. Monitoring root system width and depth changes over time via proxy measurements is a promising idea that could provide an avenue to non-destructive root system shape measurements and time course analysis of root system development.

**Figure 12:**
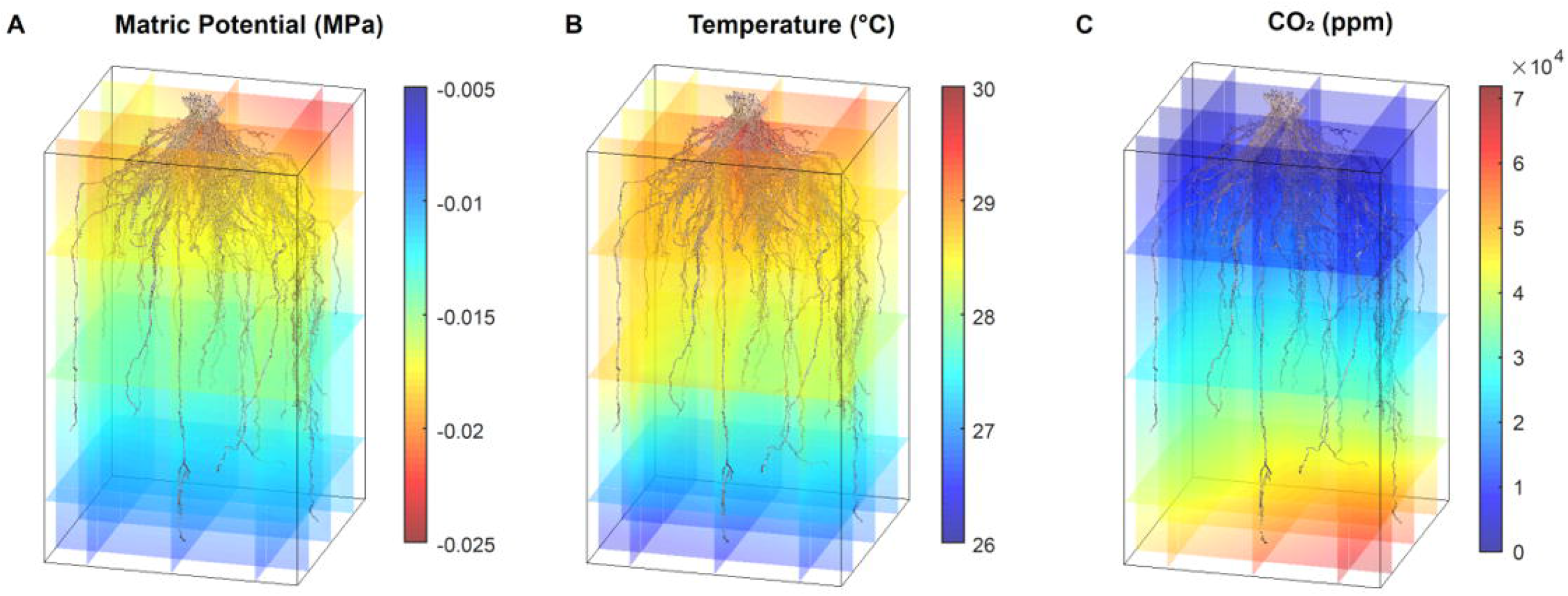
Point cloud RSA models and environmental data synthesis. Data shows one switchgrass point cloud with coaligned environmental 3D data for matric potential (A), temperature (B) and CO2 levels (C). All data are from the same time point collected between 11 am and 1 pm.

Further, direct comparisons of RSA to environmental conditions can be achieved at the cuboid level, and we have seen interesting preliminary data demonstrating a root system’s capacity to affect its surroundings. 3D interpolated environmental data was partitioned into 9×9×10” cuboids similar to the biomass measurement described in 3.2. We labeled the cuboid that contains root as ‘root cuboid’ and the cuboid that does not contain root as ‘non-root cuboid’. For every layer, we calculated the average matric potential among root cuboid and the average matric potential among non-root cuboid. The data shows that under a well-watered condition, the matric potential of root and non-root cuboids are almost identical. However, under water stressed conditions, the root cuboids are consistently wetter at every layer, with the effect being more obvious at top layers that have more water availability than the bottom layers (Fig 13). We suspect this result indicates that active root uptake is drawing water into the root occupied regions from those without, and hints at the potential to infer a coarse 3D root system architecture over time from embedded sensor data.

**Figure 13:**
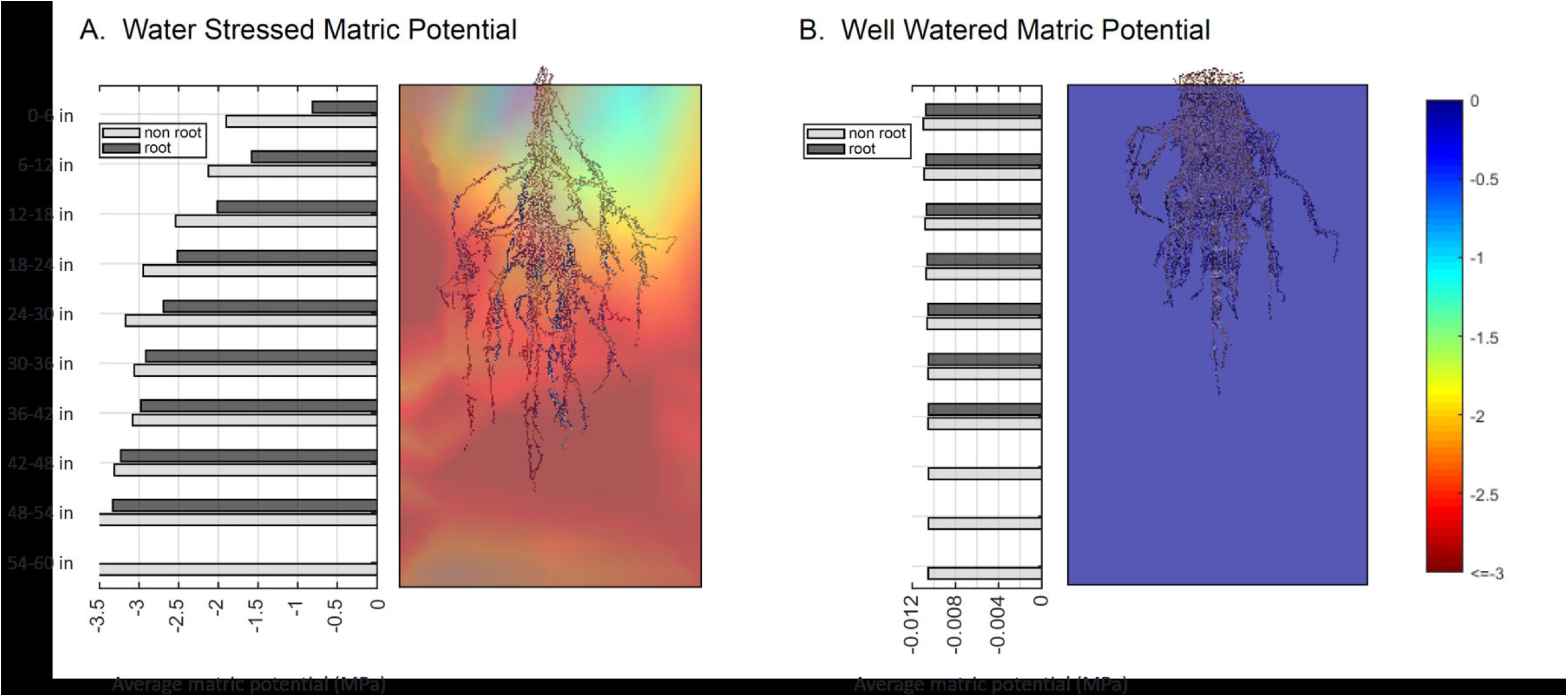
Bar plots of average matric potential for root cuboids and non-root cuboids at every layer for a water stressed sample (A) and a well-watered sample (B). Values near zero (blue) represent high water availability, while more negative values (red) denote a lower matric potential of the growth media and lower water availability.

## 4 Discussion

### 4.1 A 3D Root Mesocosm System for Integrated Environmental Sensing and Root Phenotyping

The concept of mesocosms in plant biology have been used widely to refer to a variety of experimental systems. From assessing the effects of invasive European earthworms on North American tree growth (Hale et al., 2008), to the reduction in soil-mercury emission due to soil shading by vegetation (Gustin et al., 2004), or the effects of sediment nutrition and light resources on seagrass growth and development (Short, 1987; Short et al., 1995), mesocosms are a useful intermediate between the laboratory and the field (Odum, 1984).

Root mesocosms, typically large horticultural pots or long narrow pipes form which the entire root system can be extracted, have proven useful in understanding RSA and root function of several agricultural species. For examples: it was reported that a lower number of crown roots in maize can be beneficial for nitrogen acquisition in poor nitrogen soils (Saengwilai et al., 2014), a moderate progressive drought could lead to RSA adaptations in various rice cultivars that improve performance under reduced water management practices (Hazman and Brown, 2018), some Green Revolution wheat progenitors have smaller root systems than older landraces (Waines and Ehdaie, 2007), and Chilean red clover cultivars with certain RSA traits, such as high crown root diameter and low branching index, correlate with superior persistence (Inostroza et al., 2020). However, in these studies the root systems were physically constrained during growth, leading to, at minimum, compromised estimates of root length densities and other metrics across the depth profile. To our knowledge, we report here the first system to grow large crop plants to maturity and recover unconstrained, intact root systems in their nearly-natural configuration. With the accompanying imaging and analysis plus sensor data, we have developed a new, flexible, paradigm for comprehensive subterranean analysis of root and rhizosphere biology.

### 4.2 Next Generation Mesocosms

New versions of the mesocosm are being developed to expand the scope and versatility of the technology. A large-scale version, measuring appx. 3 m wide x 6 m long x 2 m tall is being developed to more closely replicate field dynamics. In this “common-garden” or “plot-level” system, rows of plants can be placed across several frames to begin to understand multi plant dynamics. We have also added a robotic imaging system for high-throughput above-ground plant imaging. Another version is modular, with subsystems for analyzing plants with smaller root systems such as rice, wheat, and covercrops. Several can also be connected together to create fewer, but larger units as experimental needs change. These systems will accommodate a wider variety of sensors and allow access to different depths through a series of ports that allow root and rhizosphere sampling in situ. An important goal is to improve the realism of the system, and although we used artificial growth media in this study, in principle, any reconstituted soil or soil-substrate can be used. Considerations include the weight of the system and the ease and efficacy of recovering root systems intact.

### 4.3 The importance of capturing entire 3D root system architectures grown nearly unconstrained

Photogrammetry has many uses in plant biology and is a field of rapidly evolving interest (Iglhaut et al., 2019). Drone based imagery has been widely adopted as a tool to evaluate forest coverage, health and activity (Goodbody et al., 2019; Iglhaut et al., 2019; Miller et al., 2000; Mlambo et al., 2017). Similarly, terrestrial based projects such as assessments of the shape of individual trees (Bauwens et al., 2017; Gatziolis et al., 2015; Marín-Buzón et al., 2020; Wang et al., 2004) or various fruits (Feldmann and Tabb, 2022; Gené-Mola et al., 2021; Ni et al., 2021) have also become more common. A recent study has shown the power of optical reconstructions for 3D analysis of root crowns (Liu et al., 2021). However, to the authors’ knowledge, the 3D Root Mesocosms are the first system to generate 3D reconstructions of entire full grown crop root systems in nearly natural configurations, from any method.

Although the imaging of samples using photogrammetry is a low-cost process that does not require significant infrastructure, there are several challenges that still remain. Unlike other tomographic techniques, such as X-ray CT (Shao et al., 2021), photogrammetry does not resolve internal structures of the sample as the 2D images are only capturing surface features within line of sight of the 2D-photographs. This means that dense root crowns or areas of thick matted lateral roots are not resolvable. Thus, we are considering the potential to complement the photogrammetry derived point cloud with X-ray CT derived root crown reconstructions (Shao et al., 2021; Zeng et al., 2021). Additionally, photogrammetry at such a large scale can require significant computation power, dedicated software, and can currently take on the order of days to process each sample. Even considering these limitations, photogrammetry still represents a powerful tool to generate 3D models of root architecture that is flexible to image a wide array of samples, and is comparatively low-cost in relation to other tomographic methods.

The development of entire 3D root system models based on actual (non-computer-generated) plants also provides an opportunity to assess the amount of error inherent to a range of commonly utilized field-based root phenotyping methods such as soil cores, minirhizotrons, and shovelomics, which seek to estimate entire root systems from partial sampling (Pagès and Glyn Bengough, 1997; Trachsel et al., 2011; Wu et al., 2018). One idea is to generate *in silico* soil cores or minirhizotron images from the point clouds. This method could provide a sensitivity analysis for empirical sampling strategies using actual, rather than virtual (Burridge et al., 2020; Morandage et al., 2019), groundtruths. Such information could also be used as a valuable resource for improving root structure-function simulations (Kalogiros et al., 2016; Postma et al., 2017; Schnepf et al., 2018), or for the development of artificial intelligence approaches to complement missing data (Falk et al., 2020; Gaggion et al., 2021; Ruiz-Munoz et al., 2020).

### 4.4 Conclusion

The field of root system architecture phenotyping has advanced dramatically over the last few decades, from simple measurements taken with a ruler to the development of interactive virtual reality platforms. While the core complications of root phenotypic and functional analysis remain, advances along several avenues have allowed researchers to begin to analyze and visualize the subterranean dynamic complexities of root systems. We believe that, currently, coupling mesocosms and photogrammetry is a powerful way to assess the 3D structure of full grown, unconstrained root systems in their natural configurations. The methods detailed here are easily adapted to fit any size of plant and can be scaled appropriately to study concepts such as plant to plant root system interactions or planting density effects on RSA in a relatively inexpensive and easy to build manner. Further, the ease of incorporating various sensors or sampling schemes at the desired locations in the subterranean profile provides an unprecedented freedom to target specific areas of the root system to observe architectural traits and root function.

## 6 Author Contributions

Dr. Tyler Dowd developed the initial mesocosm design and lead in their physical construction, established the photogrammetry workflow to develop 3D model from the root systems, lead the selection and implementations of environmental sensors, conducted the bulk of photogrammetry and subsequent point cloud cleaning, development the measurement of biomass collection to ground truth the point clouds, conducted all subterranean CO_2_ measurements, and was the primary author of writing the manuscript.

Dr. Mao Li developed the semi-automated cleaning and segmentation of the root system point clouds, the methods of root trait extraction from the isolated root system point clouds, the visualization and modeling of 3D sensor data, the complementation of the various sensor data with the root system point clouds, and assisted in the writing of the manuscript

Dr. G. Cody Bagnall developed subsequent mesocosm designs, led the construction of mesocosms after Generation 1, developed the blueprint of the construction figures, and assisted in the writing of the manuscript.

Andrea Johnston assisted in the imaging of mesocosm root systems, the cleaning of resultant point clouds, the biomass collection to provide ground truth measurements, and assisted in the writing of the manuscript.

Dr. Christopher Topp conceived the work, oversaw and coordinated the research, and contributed to writing of the manuscript.

## 8 Abbreviations

(MRI): Magnetic Resonance Imaging
(RSA): Root System Architecture
(X-ray CT): X-ray computed tomography

## 9 Acknowledgments

The authors would like to acknowledge Dr. Amy Tabb and Michael Schoenewies for expert advice on photogrammetry, Eric Floro for preliminary experiments, Dr. Mon-Ray Shao for help in initial mesocosm design, Ben Laws for help in initial mesocosm design and fabrication of the mesocosms, Dr. David Fike for lending the Picarro system and supporting its operation, Dr. Jose Ruiz and Dr. Alina Zare for their assistance with point system skeletonization, and Dr. Samuel McInturf for the development of an R script to process Picarro Data.

## 10 Supplementary Material

Supplemental Figure 1: Interpolation of 3-dimensional environmental sensor data.

Supplemental Figure 2: Time course of shoot morphological responses of switchgrass in different growth media.

Supplemental Figure 3: Manual post-process cleaning of RSA point clouds.

Supplemental Figure 4: Comparison of the biomass and point number located in each cuboid throughout the mesocosm growth zone.

Supplemental Figure 5: Dissection of mesocosm grown root system for biomass measurements

Supplemental Figure 6: Diurnal environmental fluxes in mesocosms across nine days.

Supplemental Figure 7: Growth media effects on lateral root architecture

Supplemental Figure 8: Subterranean CO_2_ flux monitoring in mesocosms.

Supplemental Video 1: A 360-degree rotation of a photogrammetry generated point cloud of a Sorghum (BTX623) root system grown in turface under well-watered conditions.

Supplemental Video 2: A 360-degree rotation of a photogrammetry generated point cloud of a Maize (PHZ51) root system grown in turface under well-watered conditions.

Supplemental Video 3: A 360-degree rotation of a photogrammetry generated point cloud of a Switchgrass (WBC3) root system grown in turface under well-watered conditions.

Supplemental Video 4: A close up 360-degree rotation of a photogrammetry generated point cloud of a Switchgrass (WBC3) root crown grown in turface under well-watered conditions.

Supplemental Video 5: A 360-degree rotation of a photogrammetry generated point cloud of a Switchgrass (WBC3) root system grown in turface under well-stressed conditions.

Supplemental Video 6: A photogrammetry generated point cloud of a Switchgrass (WBC3) root system grown in turface under well-stressed conditions coaligned with diurnal matric potential flux data over 7 days.

Supplemental Video 7: A photogrammetry generated point cloud of a Switchgrass (WBC3) root system grown in turface under well-stressed conditions coaligned with diurnal temperature flux data over 7 days.

