## Supplementary figures and images for "Root System Architecture and Environmental Flux Analysis in Mature Crops using 3D Root Mesocosms"

### Supplementary Figure 1

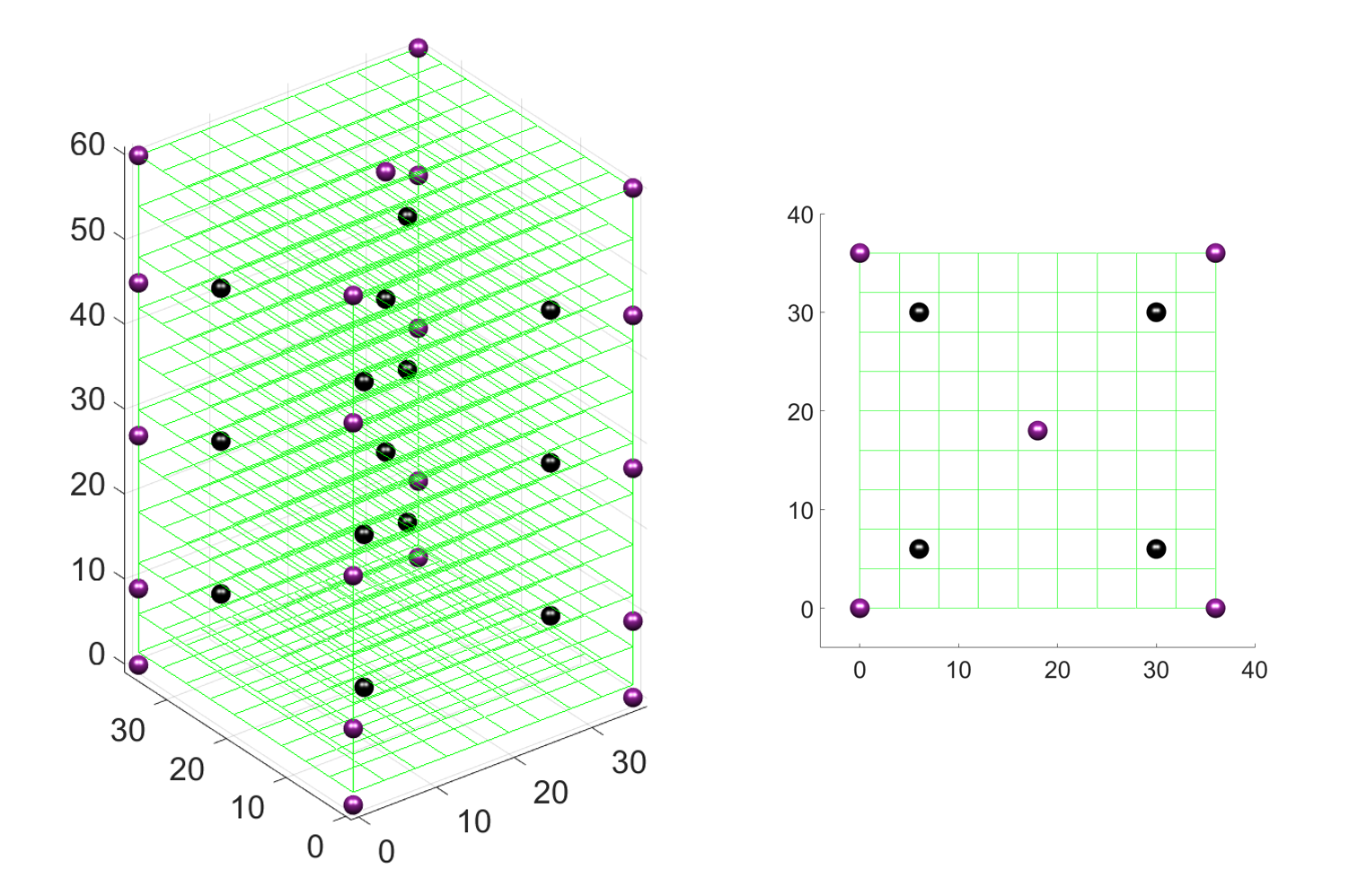

### Supplementary Figure 2

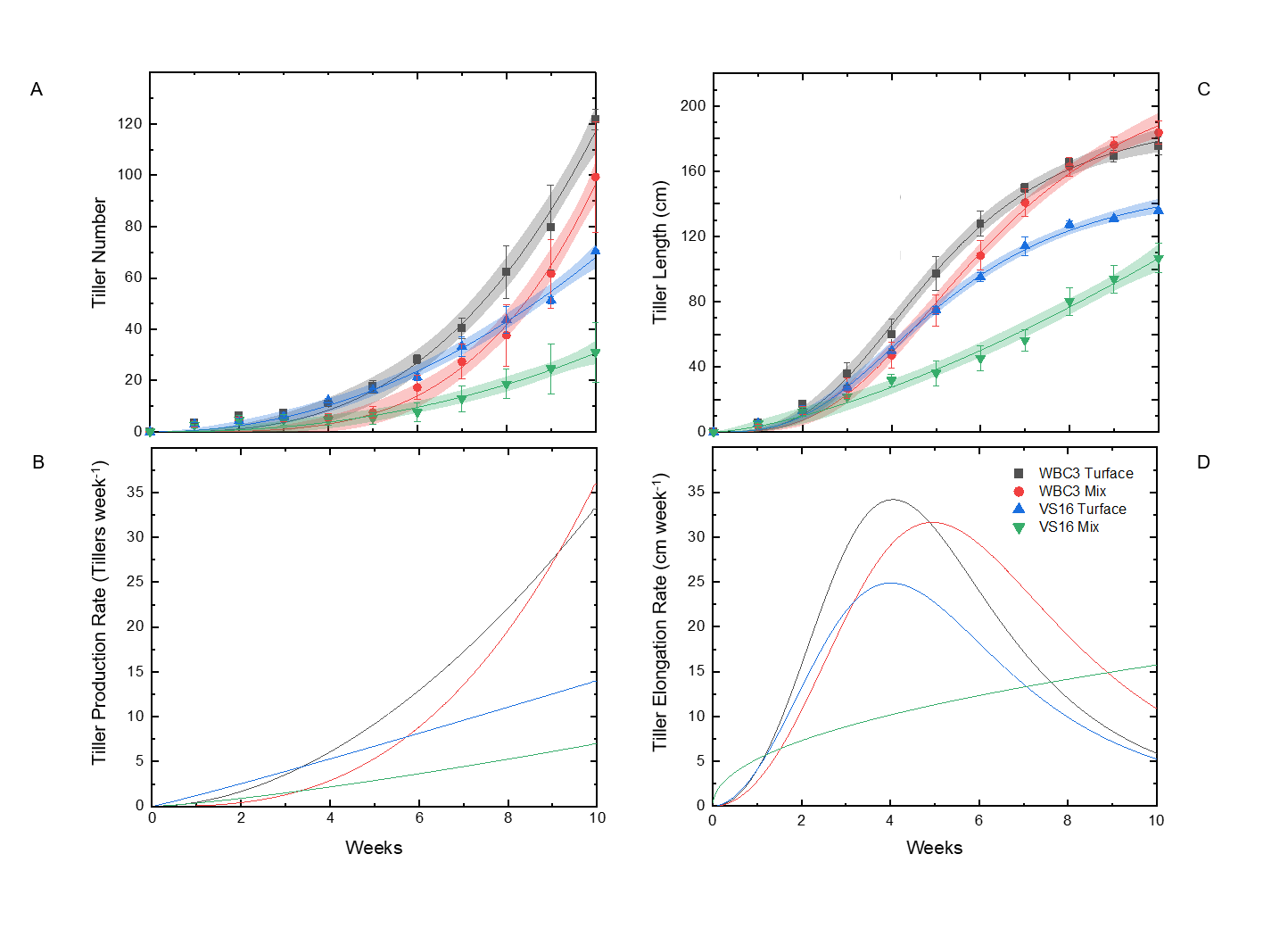

### Supplementary Figure 3

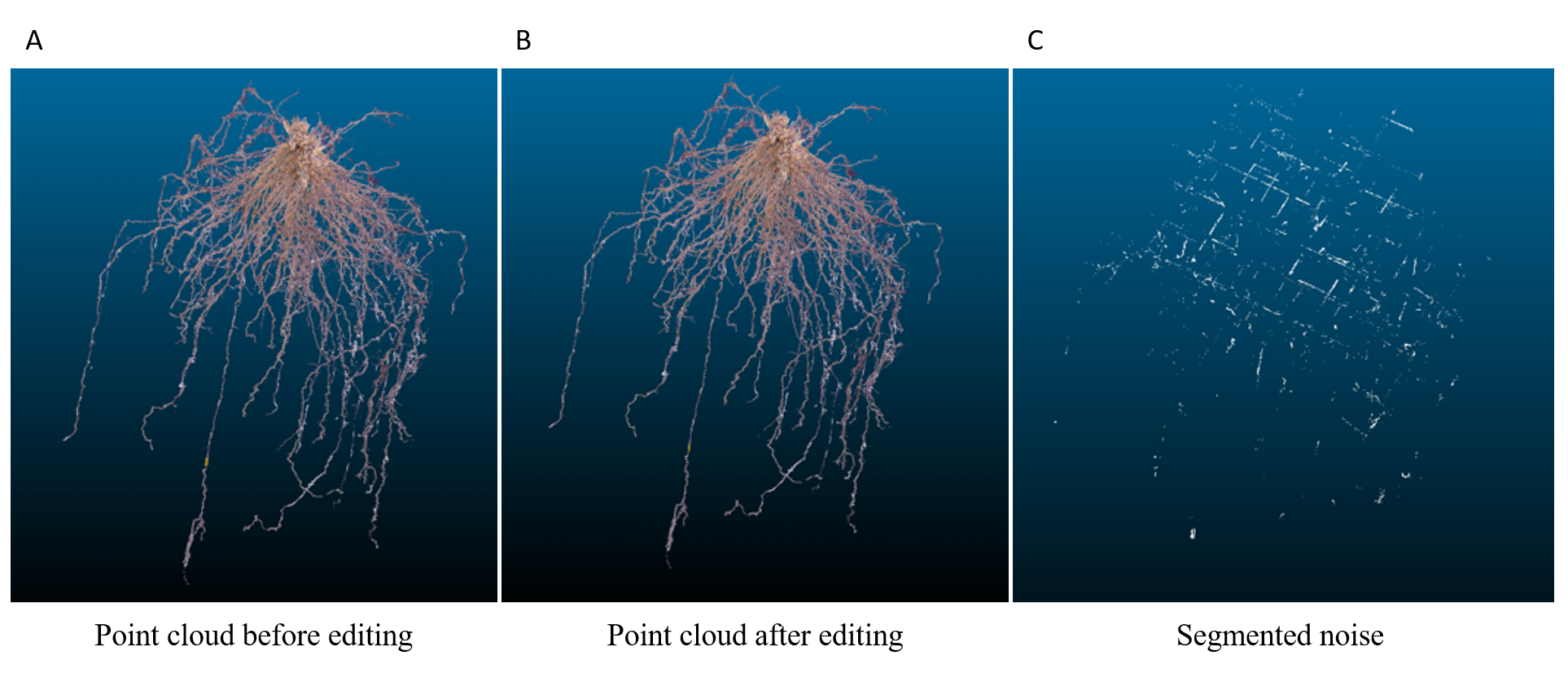

### Supplementary Figure 4

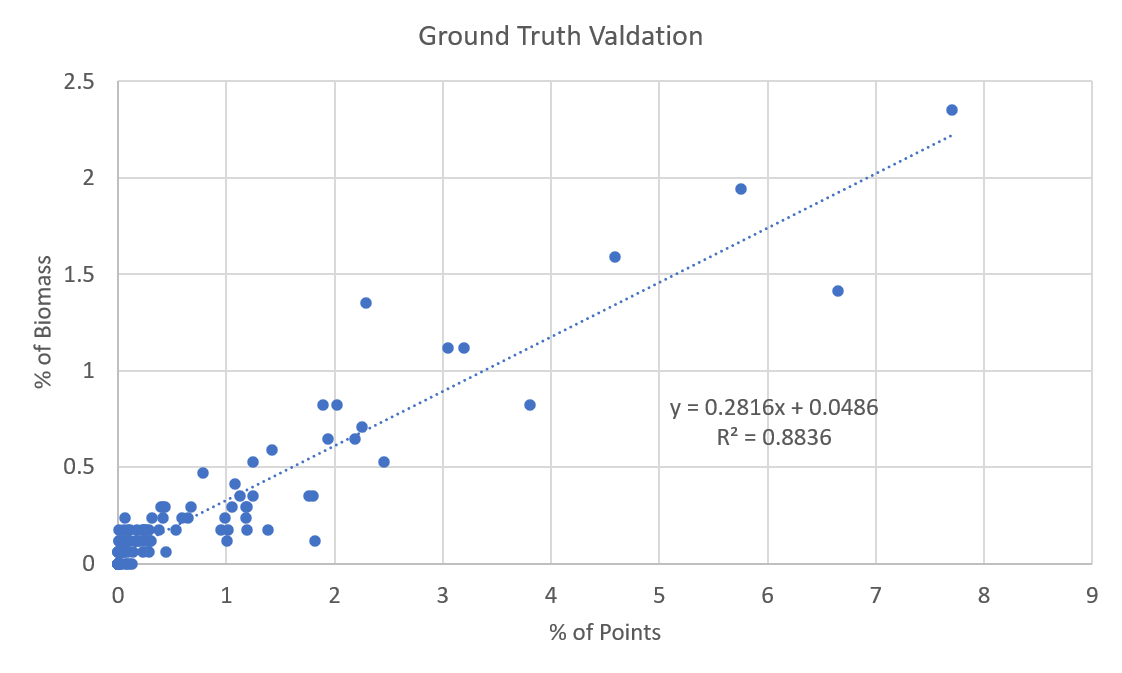

### Supplementary Figure 5

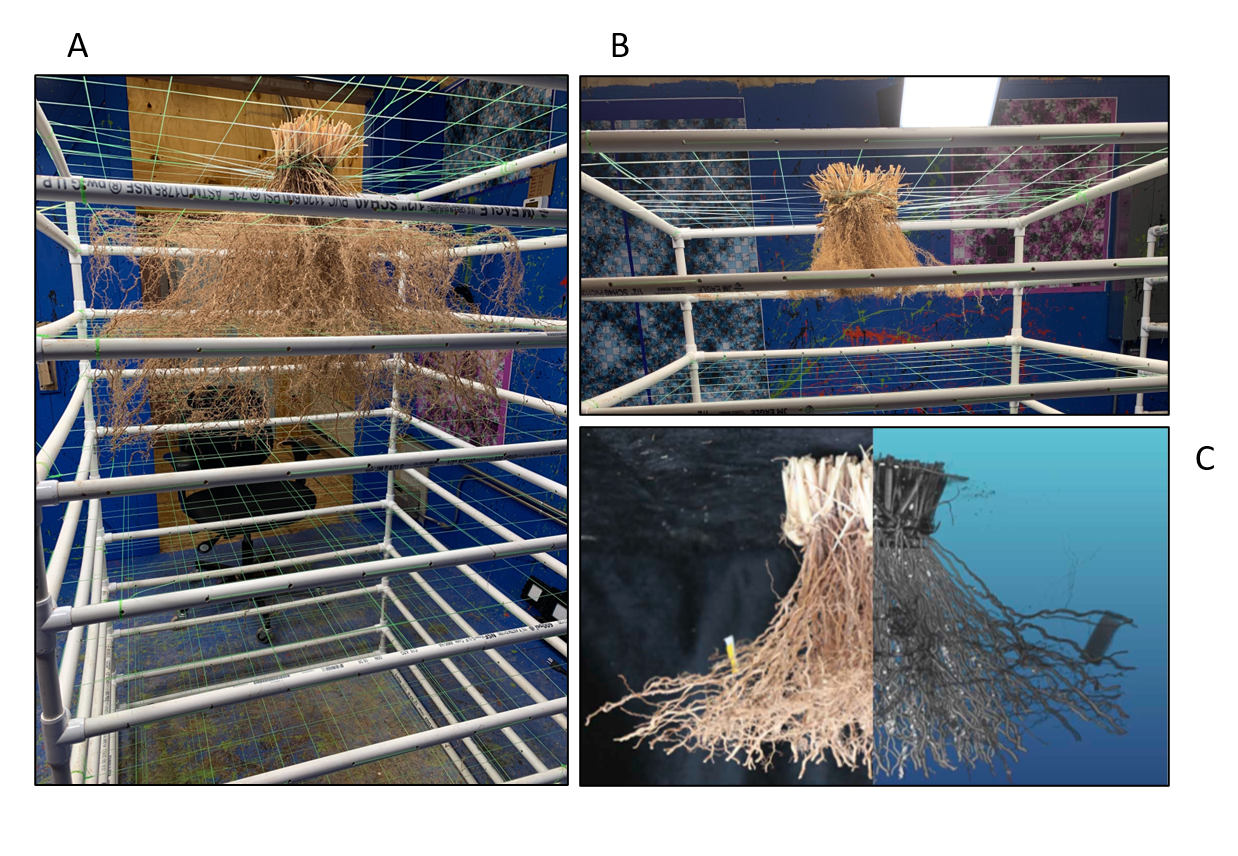

### Supplementary Figure 6

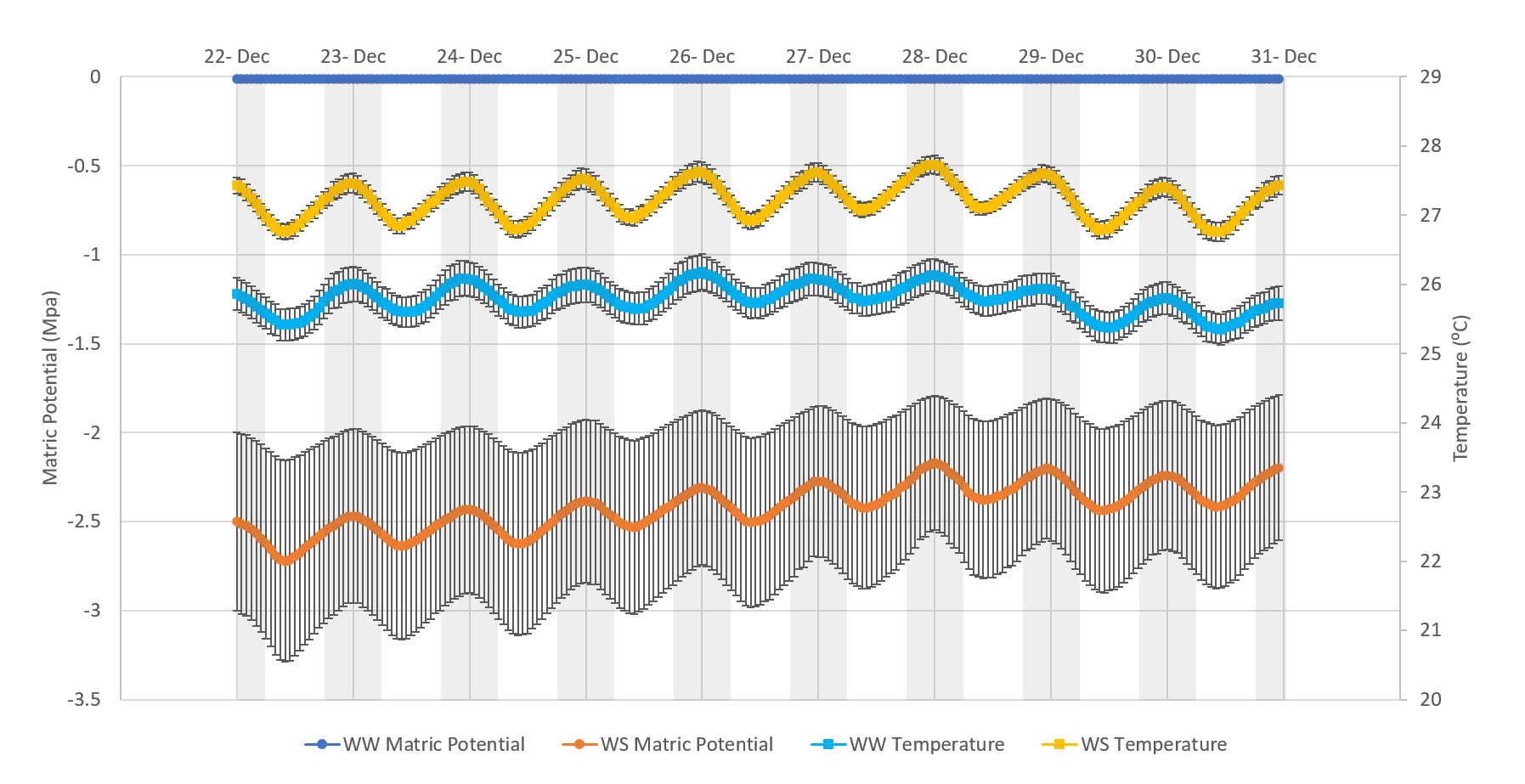

### Supplementary Figure 7

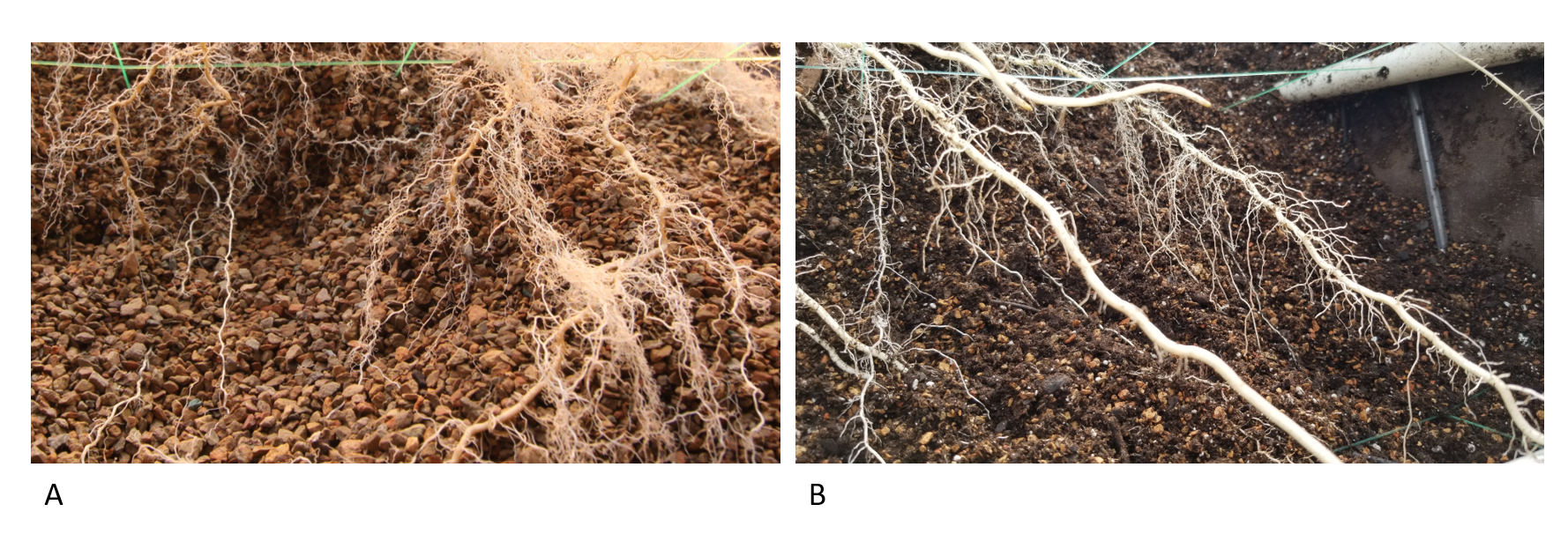

### Supplementary Figure 8

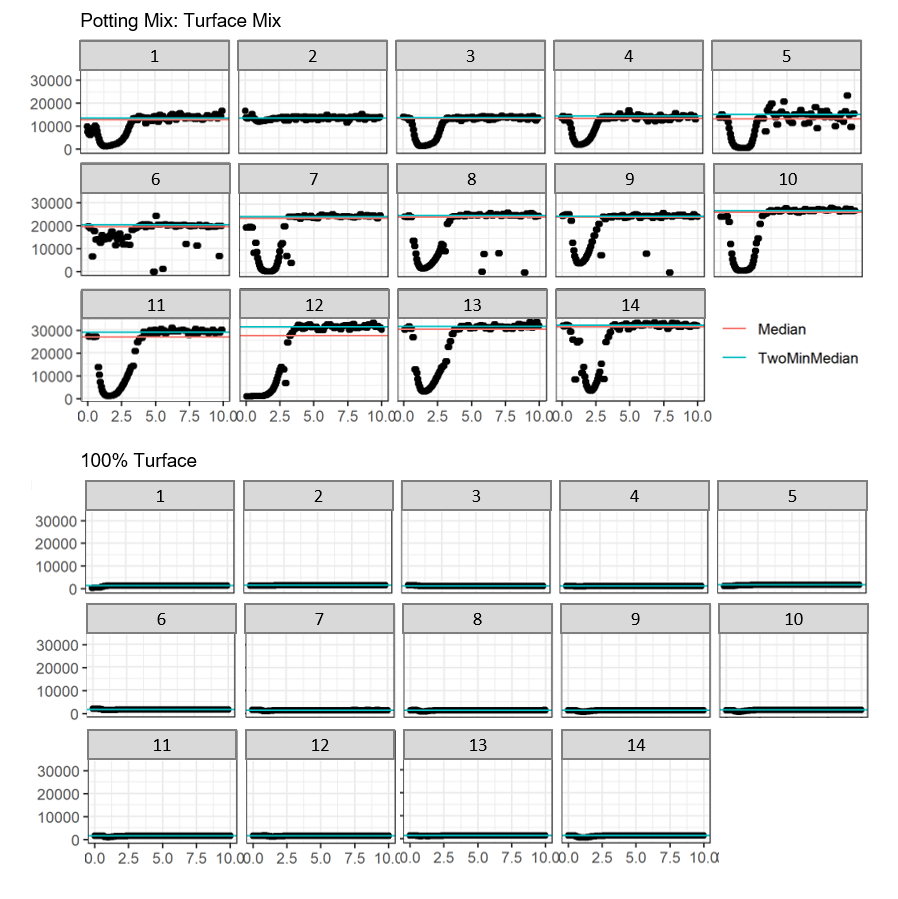
